# STING Nuclear Partners Contribute to Innate Immune Signalling Responses

**DOI:** 10.1101/2020.12.21.423744

**Authors:** Charles R. Dixon, Poonam Malik, Jose I. de las Heras, Natalia Saiz-Ros, Flavia de Lima Alves, Mark Tingey, Eleanor Gaunt, A. Christine Richardson, David A. Kelly, Martin W. Goldberg, Greg J. Towers, Weidong Yang, Juri Rappsilber, Paul Digard, Eric C. Schirmer

**Author notes:** Labstat International Inc., 262 Manitou Dr, Kitchener, Ontario, N2C 1L3, Canada (Flavia de Lima Alves). Correspondence to: Eric Schirmer, Institute of Cell Biology, University of Edinburgh, Kings Buildings, Swann 5.22, Mayfield Road, Edinburgh EH9 3BF, UK.

## Abstract

STING and cGAS initiate innate immune responses (IIR) by recognizing cytoplasmic pathogen dsDNA and activating signaling cascades from the ER; however, another less investigated pool of STING resides in the nuclear envelope. We find that STING in the inner nuclear membrane increases mobility and changes localization upon IIR activation both from dsDNA and poly(I:C) stimuli. We next identified nuclear partners of STING from isolated nuclear envelopes. These include several known nuclear membrane proteins, bromodomain and epigenetic enzymes, and RNA- or DNA-binding proteins. Strikingly, 17 of these DNA and RNA-binding STING partners are known to bind direct partners of the IRF3/7 transcription factors that are central drivers of IIR. We find that several of these STING partners —SYNCRIP, Men1, Ddx5, snRNP70, RPS27a, Aatf— can contribute to IIR activation and SYNCRIP can moreover protect against influenza A virus infection. These data suggest that the many roles identified for STING likely reflect its interactions with multiple RNA and DNA-binding proteins that also function in IIR.

## Introduction

The innate immune response (IIR) is the first line of defense against pathogens, recognizing molecular patterns in infected cells such as the presence of cytoplasmic DNA or dsRNA^1, 2^. STING (STimulator of INterferon Genes) (also called MITA, ERIS, MPYS, NET23, and TMEM173) is the essential adaptor protein in innate immune signaling cascades triggered by cytosolic DNA. Cyclic GMP-AMP synthase (cGAS) senses cytoplasmic dsDNA and catalyzes the synthesis of a second messenger, cGAMP, which binds to and activates dimeric STING at the ER, activating IIR signaling cascades that stimulate IRF3/7 transcription factors to activate IIR genes such as type I interferons (IFN)^3, 4, 5, 6, 7, 8, 9^. The cGAS-STING pathway for IIR triggered by cytoplasmic dsDNA has been well characterized in molecular detail; however, STING clearly has other important functions. As well as restricting the proliferation of DNA viruses through type-I IFN induction, STING also restricts the proliferation of some RNA viruses^10, 11, 12^, although the mechanisms through which it does so remain to be fully elucidated. Although STING does not directly bind to RNA, the replication of multiple positive- and negative-sense RNA viruses is enhanced in the absence of STING^6, 7, 11, 12, 13, 14, 15, 16, 17, 18, 19, 20, 21^. Several reports have argued that STING is not involved in interferon activation in response to foreign RNA^7, 13, 22^; so how it acts against RNA viruses is less clear than its counteraction of DNA viruses through the induction of type I IFN. Moreover, other IIR roles for STING have been identified in pro-apoptotic signaling with MHC II from the plasma membrane^23^, the induction of autophagy^24^, and in NF-κB activation downstream of DNA damage^25^. The many distinct functions and localizations reported for STING in IIR make it difficult to distinguish direct from downstream signaling effects.

Though originally introduced in a proteomics study of the nuclear envelope (NE) as NET23^26^, potential nuclear roles for STING have been largely unaddressed. A significant pool of STING was confirmed in the NE that was partly lost in the absence of lamin A^27^, suggesting it can reside in the inner nuclear membrane (INM); however, super resolution microscopy employed in that study suggested it was limited to the outer nuclear membrane (ONM). A subsequent study showing a role for STING in promoting chromatin compaction further suggested a function inside the nucleus^28^, though without demonstration of a pool inside the nucleus.

The potential importance to IIR of a pool of STING inside the nucleus has been enhanced greatly by recent reports of cGAS, its upstream partner in cytoplasmic dsDNA sensing, in the nucleus^29, 30, 31, 32, 33^. cGAS directly binds DNA and so a long-standing question in the field was how it was prevented from binding and being activated by chromosomal DNA. Recent studies have shown that in fact a large portion of cGAS is in the nucleus bound to chromosomes^3, 29, 31, 34^, so it is critical to keep this pool from activating IIR. At the same time, this also raises the possibility that nuclear cGAS could sense pathogen nucleic acids inside the nucleus as well as in the cytoplasm and thus activate IIR from inside the nucleus in which case nuclear STING could be required for such signaling responses.

Here we confirm inner nuclear membrane residence for endogenous STING and show that tagged nuclear STING can redistribute from the nucleus to the ER upon treatment with dsDNA or, surprisingly, the dsRNA mimetic poly(I:C). We further show that STING dynamics in the nuclear envelope increase with both DNA and RNA triggered immune responses. Moreover, we identify NE partners for STING, which are enriched for RNA and DNA binding proteins, and testing several of these partners indicates that they can contribute to IIR activation and one of the partners identified, SYNCRIP, can moreover protect against influenza and RNA virus infection.

## Results

### STING targets to the inner nuclear membrane

An earlier attempt to determine if the NE pool of STING was in the outer (ONM) or inner (INM) nuclear membrane was inconclusive^27^. Therefore, we used structured illumination (OMX) super resolution microscopy that can distinguish INM proteins from ONM proteins by their being in the same plane with respectively nuclear basket or cytoplasmic filament proteins of the nuclear pore complexes (NPCs) that are separated by ~100 nm (Fig. 1a)^35^. Analyzing several cells on the same coverslip revealed STING-GFP to be in the ONM of some cells and the inner nuclear membrane (INM) of other cells (Fig. 1b). The finding of some cells with STING in the ONM and others in the INM was striking, as most NE membrane proteins tested with this method were unambiguously resident in either the INM or ONM^36, 37^. This suggested STING may redistribute into different nuclear locations under certain cell-specific conditions.

**Fig. 1.**
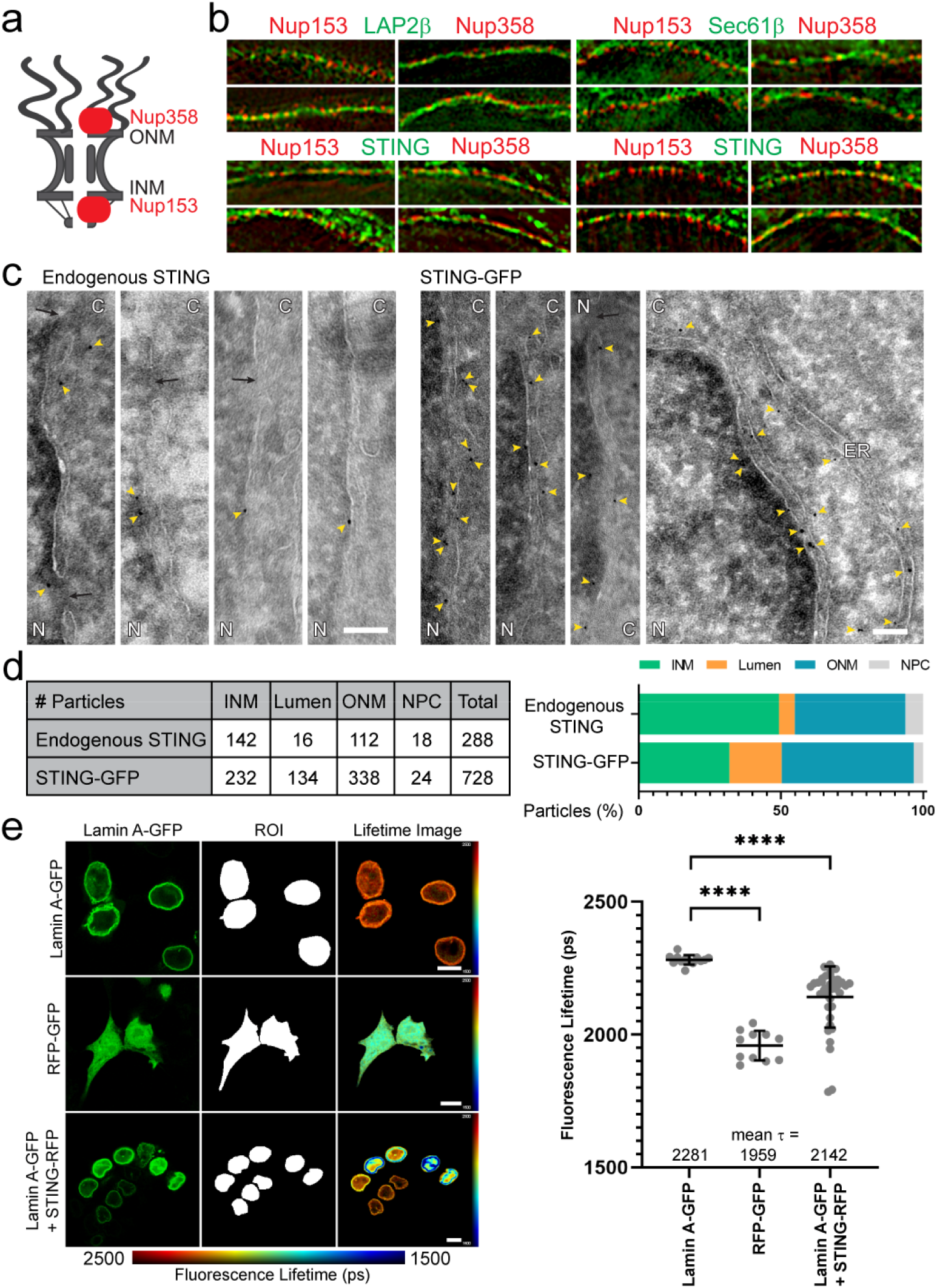
STING inner nuclear membrane localisation. **a** Schematic of nuclear pore complex (NPC) indicating location of Nup358 and Nup153 in the NPC. **b** With structured illumination super-resolution (OMX) microscopy, proteins lining up in the same plane as Nup153 indicates localisation in the inner nuclear membrane while proteins lining up in the same plane as Nup358 indicates localisation in the outer nuclear membrane. Upper panels controls: LAP2β is known to be in the inner and Sec61β in the outer nuclear membrane. Lower panels: STING is in the inner nuclear membrane in some cells and the outer nuclear membrane in others. Scale bar, 5 μm. **c** Immunogold electron microscopy for endogenous STING confirms its inner nuclear membrane localisation (Endogenous STING panels, antibody specificity confirmed in Supplemental Figure 1 and 2) with particles also observed in the outer nuclear membrane and ER. A much higher number of particles could be observed per image for exogenously expressed STING tagged with GFP that yielded a similar distribution. N, nucleus; C, cytoplasm; yellow arrowheads, immunogold particles; black arrows, NPCs; scale bar, 100 nm. **d** Quantification of the larger volume of data represented in Figure 1c. The apparent increase of particles in the NE lumen for STING-GFP likely reflects enhancement of sectioning artefacts due to the size of the tag. **e** FRET-FLIM indicates an interaction between STING and the lamin A polymer that lines the inner nuclear membrane. Representative images are shown for the lamin A-GFP alone (negative control), a tandem GFP-RFP construct (positive control), and the lamin A-GFP:STING-RFP pairing. Blue indicates a reduction in GFP fluorescence lifetime due to the transfer of photons to the acceptor RFP molecules. Quantification of averaged τ values for the fluorescence lifetime of the donor GFP signal in picoseconds revealed a significant transfer of energy from lamin A to STING, indicative of their interacting. Ordinary one-way ANOVA with Dunnett’s multiple comparison test, **** p ≤ 0.0001. Scale bar, 10 μm.

As the super resolution approach used STING fused to GFP, we also analyzed endogenous STING within the NE using immunogold electron microscopy (Fig. 1c, left images). Similar numbers of gold particles were observed at the INM (142) as at the ONM (112). Specificity of immunogold labelling against endogenous STING was confirmed by the absence of gold particles in samples stained only with secondary antibodies (Supplementary Fig. S1a). To further determine whether the GFP fusion interfered with this distribution, a cell line stably expressing STING-GFP was also analyzed by immunogold electron microscopy (Fig. 1c, right images). The distribution of particles between inner and outer nuclear membranes was not notably altered by the tag (Fig. 1d). A larger proportion of particles could not be clearly ascribed to either INM or ONM as they were detected in the lumen between the two membranes of the NE. This is likely a result of an increased distance between gold particle and protein due to the addition of the C-terminal GFP tag combined with sectioning resulting in STING-GFP residing within the INM or ONM appearing as luminal by immunogold EM.

Finally, we confirmed the inner nuclear membrane pool of STING by the ability of a C-terminally tagged STING-RFP construct to accept photons from lamin A-GFP through Förster resonance energy transfer by fluorescence lifetime imaging (FRET-FLIM). The lifetime of the activated lamin A-GFP fluorescence was reduced when cells also expressed STING-RFP as a photon acceptor (Fig. 1e). On average this INM pool of STING dropped the mean lifetime (τ) of lamin A-GFP from 2.281 to 2.142 ns. Expected targeting of STING-RFP to the ER/NE was confirmed by confocal microscopy (Supplementary Fig. S1b).

### STING dynamics are altered upon stimulation of IIR

One possible explanation for the finding of STING-GFP in the inner nuclear membrane of some cells and not in others in Figure 1b is that its localization might be altered by activation of IIR, especially as the transient transfections used for most of these experiments could have stimulated IIR in a subpopulation of transfected cells due to sensing plasmid DNA. Although nuclear redistribution of STING during IIR has not been investigated, STING accumulates in perinuclear aggregates, with a visible decrease in nuclear localization, upon IIR activation with dsDNA but not dsRNA^7, 13^ (see also Supplementary Fig. S2). This implies that, at least in response to DNA immune stimulation, STING at the INM must translocate to peri-nuclear foci, as for STING localized in the peripheral ER. Measurement of STING dynamics required using fluorescently tagged fusion proteins, therefore, we first compared the redistribution of STING in a stable STING-GFP cell line with that of endogenous STING using two different antibodies and fixations and stimulating IIR with either plasmid DNA or poly(I:C), finding a similar redistribution pattern for the dsDNA and a similar lack of visible redistribution for the dsRNA mimic, poly(I:C) (Supplemental Fig. S2a-d).

We hypothesized that if STING activation promotes shuttling between the nucleus and cytoplasm, its mobility measured by fluorescence recovery after photobleaching (FRAP) should change upon IIR induction. To avoid unintentional IIR induction due to transfected plasmid DNA, STING-GFP or a control nuclear envelope transmembrane protein (NET) fused to GFP (NET55) were stably expressed in HT1080 cells. Induction of IIR by infection with herpes simplex virus type 1 (HSV-1) visibly increased the speed of fluorescence recovery (Fig. 2a), reducing the t½ for STING in the NE by ~⅓ from 11.1 to 6.7 s (Fig. 2b,c). In contrast, the t½ of control NET55 was unaffected. This most likely indicates an increase in STING shuttling upon IIR activation, because NE FRAP has been shown to principally measure translocation through the peripheral channels of the NPC^38^. Interestingly, STING was not observed to accumulate in peri-nuclear foci as occurs with dsDNA stimulation. However, this is consistent with reports that HSV-1 inhibits STING activation and can prevent translocation of STING to the Golgi apparatus^39, 40, 41^. Suprisingly, STING mobility in the NE was also increased by stimulation with poly(I:C) (Fig. 2d), dropping the t½ from 13.3 s for the control in that experiment to 7.59 s (Fig. 2e). This was unexpected because the STING perinuclear accumulation is known to only occur in response to DNA stimuli and not to poly(I:C), suggesting this increased mobility of STING for both stimuli is not related to the canonical pathway.

**Fig. 2.**
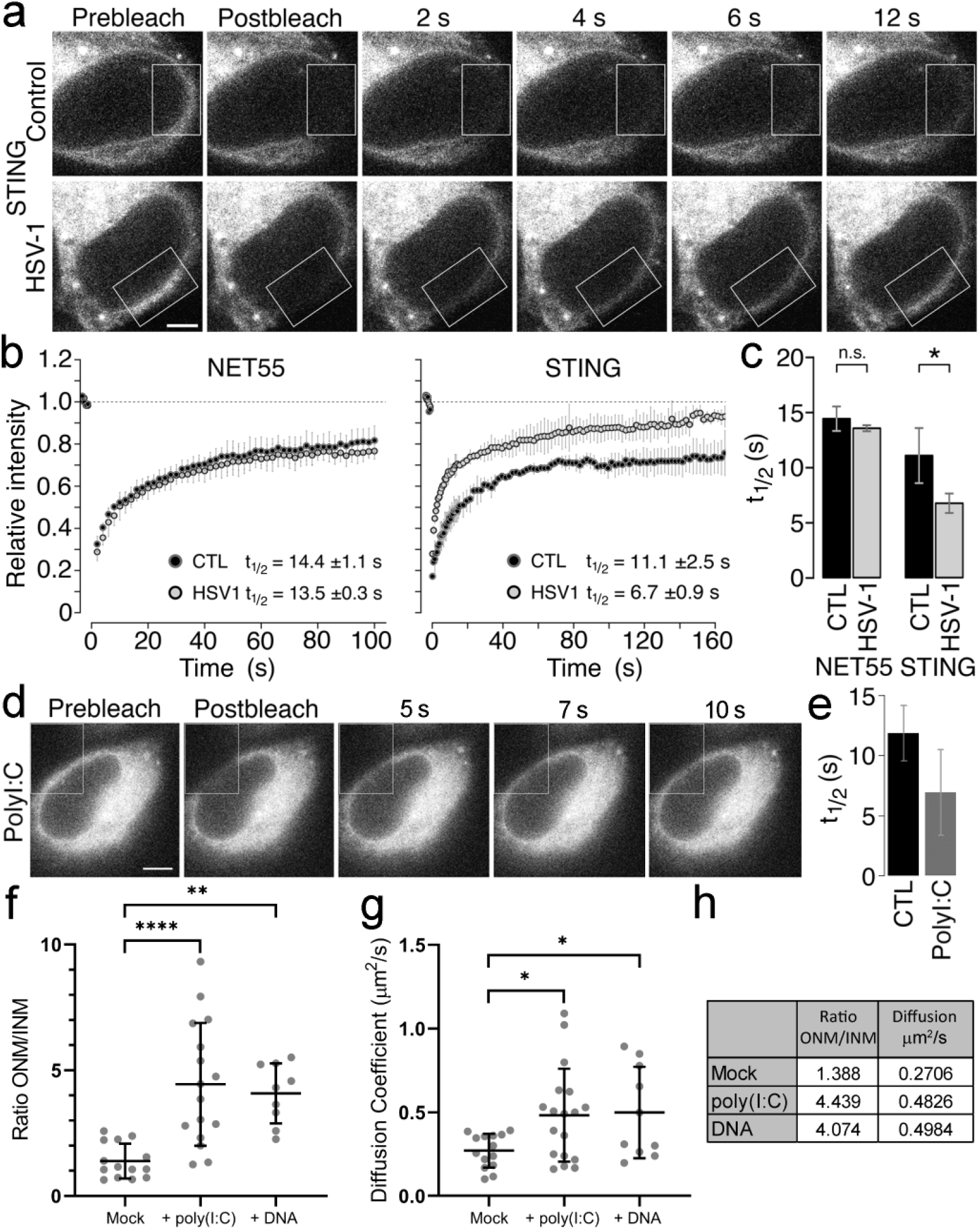
STING nuclear membrane mobility increases upon IIR activation. **a** FRAP of STING-GFP in control (mock-infected) and HSV-1 infected cells (2 hpi), photobleaching an area within the white outlined box. Scale bar, 5 μm. **b** Fluorescence recovery curves from three replicate experiments as in *a*. Another NE protein, NET55, is shown as a control that does not change its dynamics with HSV-1 infection. CTL, control; HSV1, HSV-1 infected. **c** Bar plot comparing the average half recovery times (t_1/2_) between the control and HSV-1 infected cells (student’s T test, * p ≤ 0.05). **d** FRAP of STING-GFP in cells 2 h after poly(I:C)-treatment, photobleaching an area within the white outlined box. Scale bar = 5 μm. **e** Bar plot comparing the average half recovery times (t_1/2_) between the untreated control and the poly(I:C) treated cells. **f** SPEED microscopy on control and poly(I:C) or dsDNA transfected cells expressing STING-GFP revealed a redistribution from the inner nuclear membrane to the outer nuclear membrane/ER compartment. The ratio of particles in the outer nuclear membrane (ONM) over the inner nuclear membrane (INM) is plotted. Statistics used ordinary one-way ANOVA with Dunnett’s multiple comparisons test, ** p ≤ 0.01, **** p ≤ 0.0001. **g** Measurement of mobility in the form of the diffusion coefficient measured in the same SPEED microscopy experiments as in *f*revealed also increased mobility induced by polyI:C or dsDNA. Statistics used ordinary one-way ANOVA with Dunnett’s multiple comparisons test, * p ≤ 0.05. **h** Table of summary data from *f* and *g* displaying mean values.

We next turned to a different super resolution approach, Single-Molecule Fluorescence Recovery After Photobleaching (smFRAP) microscopy, that enables the tracking of individual NETs as they diffuse along the INM and ONM of the NE^42, 43, 44^. The nuclear pool of STING-GFP redistributed out of the nucleus upon stimulation of the cells with poly(I:C) or dsDNA (Fig. 2f). A similar number of STING-GFP molecules were in the inner versus outer nuclear membranes (as was observed by immunogold electron microscopy) in unstimulated cells, while the ratio of outer nuclear membrane to inner nuclear membrane signals more than doubled in the poly(I:C) and dsDNA (Fig. 2f and h) stimulated cells. Measuring the diffusion coefficient of this mobile STING revealed a near doubling of the speed of particles from 0.27 to 0.48 μm^2^/s in the poly(I:C) treated cells and to 0.49 μm^2^/s in dsDNA treated cells (Fig. 2g and h).

### Many STING NE partners are nucleotide binding proteins

Having established that STING is present in the INM, we sought to investigate its role in the nucleus by identifying NE-specific partners that may have been missed in earlier studies because STING’s association with the intermediate filament lamin polymer^27^ likely renders this pool highly insoluble. In order to maintain interactions with potential NE partner proteins when subsequently disrupting STING’s strong association with the INM and nuclear lamina, NEs were first isolated from HEK293T cells transiently expressing STING-GFP and then treated with a reversible cross-linker. Cross-linking chased STING-GFP into complexes between 130 and 300 kDa that could be reverted to the expected 70 kDa upon reversal of the cross-linking with DTT (Fig. 3a and b). Cross-linked NEs were fragmented by sonication, immunoprecipitated (IP’d), cross-links reversed, and putative partners identified by tandem mass spectrometry (Table S1). The proteins that co-IP’d in the STING-crosslinked NEs were weighted for likely abundance based on spectral counts and plotted based on Gene Ontology (GO)-biological process terms. This revealed an enrichment in proteins with GO-terms for chromatin/chromosome organization and RNA/DNA binding compared to their representation among all proteins encoded by the genome (Fig. 3c). Plotting the normalized spectral abundance in the STING-GFP sample compared to mock transfected cells revealed the most abundant of the enriched co-IP proteins to be histone H1 variants (Fig. 3d) followed (Fig. 3e) by a mixture of known NE proteins (e.g. Lamin A, LAP2), nucleotide-binding proteins (e.g. snRNP70, UBTF, RPS27a), and bromodomain proteins (e.g. Brd2, Brd3, Rbmx) that could mediate the reported STING function in chromatin compaction^28^ and other epigenetic changes associated with IIR^45^ (Table 1). Many proteins in all these categories bind DNA/RNA and nearly half of all STING partners identified are listed as nucleotide-binding proteins (Fig. 3f). Strikingly, although some known STING interactors were identified in the NE-STING proteome (DDX41^46^ and CCDC47^47^) many well-known interactors such as TBK1 and MAVS were not found, suggesting that the NE-STING proteome differs significantly from that of STING localized in the ER.

**Fig. 3.**
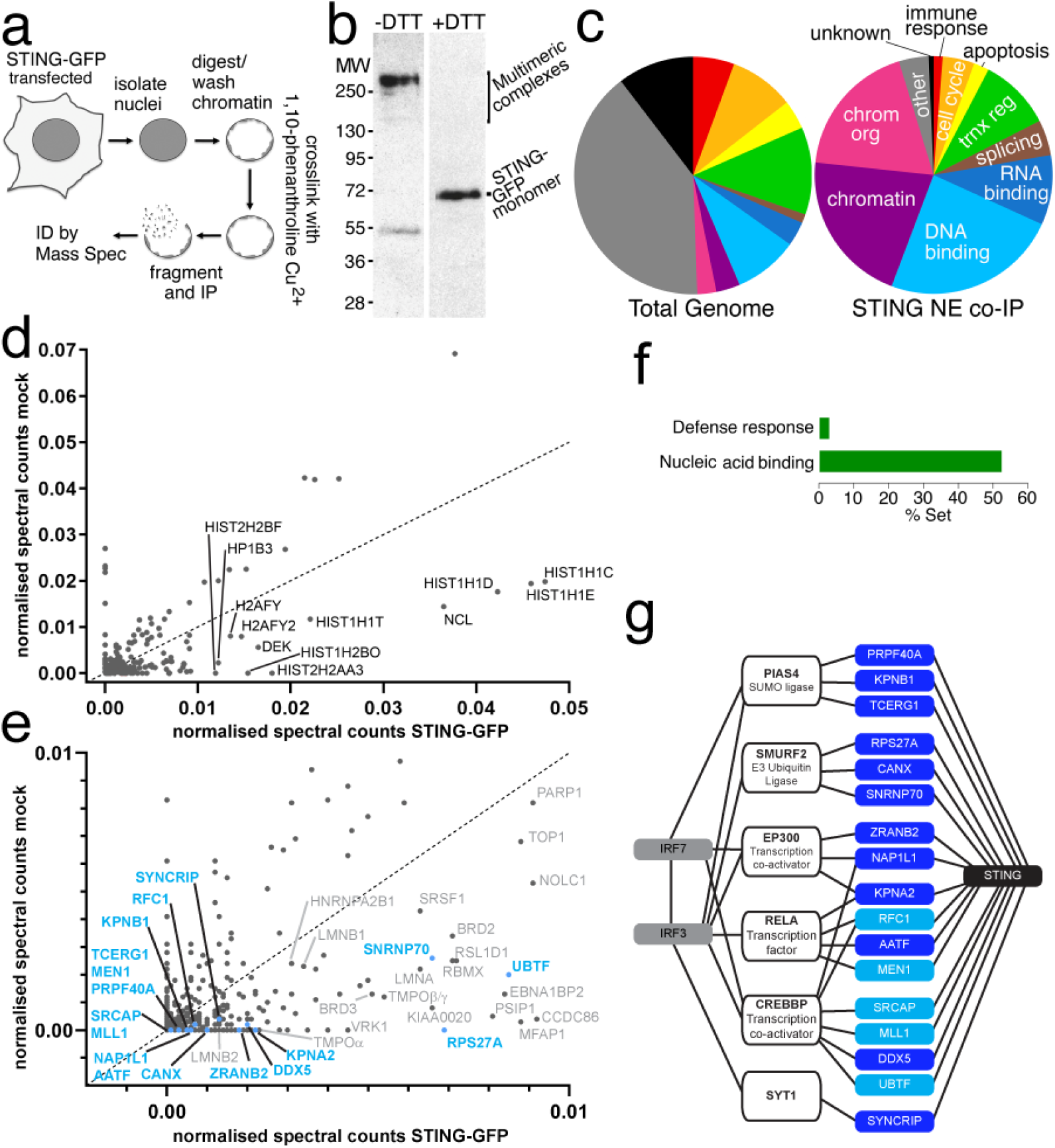
Many proteins identified by STING NE co-IP have nucleotide-binding functions. **a** Schematic of reversible-crosslinking approach. NEs were isolated from HEK293T cells expressing STING-GFP or mock transfected cells. The NEs were crosslinked with orthophenanthroline copper, fragmented by sonication and STING-GFP crosslinked proteins recovered by immunoprecipitation with GFP antibodies. The crosslinking was reversed to release these other proteins and their identity determined by mass spectrometry. **b** Crosslinking of NEs with orthophenanthroline copper chases most STING-GFP to multimeric species >200 kDa while a smaller portion appears at 55 kDa presumably due to intramolecular crosslinks. DTT-induced reversal of crosslinking restores all STING-GFP to its expected molecular weight at ~69 kDa. **c** Gene ontology (GO) biological process classification for STING putative NE partners identified by mass spectrometry of crosslinking NE co-IP material. The representation of the GO-terms by number of genes in the total human genome is shown on the left while on the right are the terms as represented in the STING co-IP material with weighting based on the number of spectra recovered from each protein. **d** putative STING NE partners plotted by normalized spectral abundance and enrichment in STING-GFP samples versus control samples. Nearly all of the most abundant partners were histone H1 variants. **e** Plotting the same data in *d,* but only up to 0.01 for normalized spectral abundance. The position of the proteins indicated by the analysis in panel *g* is highlighted in blue. **f** Bar graph showing the representation within the set of putative STING NE partners of all GO-terms associated with host defense responses or nucleic acid binding. **g** Known interacting proteins for the putative STING NE partners identified from the reversibly crosslinked NEs were searched for using the HPRD interactome database. 17 of the putative STING NE partners (blue) had reported interactions with 6 proteins (white boxes) reported to bind IRF3/7 transcription factors (grey) central to IIR activation.

**Table 1.** Functional groups of STING highest abundance NE interactors based on raw spectral counts

| Epigenetic | s | NET/NE | s | Histone | s | RNA | s | Other | s |
| --- | --- | --- | --- | --- | --- | --- | --- | --- | --- |
| HP1B3 | 77 | LMNA | 48 | HIST1H1C | 104 | PSIP1 | 50 | NCL | 287 |
| BRD2 | 64 | TMPOβ | 28 | HIST1H1E | 103 | SNRNP70 | 35 | UBF1 | 78 |
| KIAA0020 | 50 | TMPOα | 18 | HIST1H1D | 97 | RBM28 | 33 | DEK | 73 |
| BRD3 | 42 | CKAP4 | 15 | HIST2H2AA3 | 26 | HNRNPG | 31 | MFAP1 | 47 |
| BAZ2A | 28 | KPNA2 | 13 | HIST1H2BO | 22 | EBNA1BP2 | 30 | CD11B | 46 |
| MECP2 | 19 |  |  | HIST2H2BF | 17 | RRP1B | 24 | RL1D1 | 41 |
| HELLS | 16 |  |  |  |  | HNRNPR | 23 | CCDC86 | 38 |
| RSF1 | 12 |  |  |  |  | HNRNPL | 18 | NOP2 | 23 |
|  |  |  |  |  |  | PAF1 | 18 | VRK1 | 21 |
|  |  |  |  |  |  | HNRNPK | 17 | ILF3 | 16 |
|  |  |  |  |  |  | RRMJ3 | 15 | KIF22 | 16 |
|  |  |  |  |  |  | RBMXL1 | 15 | HDGR2 | 15 |
|  |  |  |  |  |  | DDX5 | 14 | NOP58 | 14 |
|  |  |  |  |  |  | SRSF2 | 13 | GTF2I | 14 |
|  |  |  |  |  |  | RPS27A | 13 | UHRF1 | 13 |
|  |  |  |  |  |  |  |  | HSPA1A | 13 |

NOTE: restricted to those with >3x more spectra (s) in STING sample than in mock sample

Downstream of STING ER/Golgi functions, IRF3/7 transcription factors induce IFN and other IIR genes in the nucleus. Therefore, we wondered if some of these STING NE coIP partners have known interactions with IRF3/7 and may modulate immune signaling cascades. Accordingly we searched the HPRD interactome database^48^ using Cytoscape^49^, finding that IRF3/7 had no known direct interactions with any of the putative STING partners. However, six known direct IRF3/7-binding partners interact directly with 17 of the proteins identified in the STING-NE co-IP (Fig. 3g). Of these, 12 are RNA-binding proteins (dark blue). The rest, as well as some of the RNA binding proteins, have also been reported to bind DNA. Although these proteins have not been previously shown to affect IRF3/7 transcriptional responses in IIR, several interact with viral proteins and affect viral replication, so may contribute to host cell IIR. For example, DDX5 is bound by the N(pro) protease of pestivirus^50, 51^ and may inhibit hepatitis C virus replication^52^ and vesicular stomatitis virus triggered IFNβ induction^53^, although it appears to be a positive regulator of HIV-1^54^, and Japanese encephalitis virus (JEV)^55^ among others^56^. Meanwhile, the hepatitis B virus HBx protein alters the intracellular distribution of RPS27a^57^, AATF is specifically targeted by an HIV-encoded miRNA^58^, and SYNCRIP is involved in hepatitis C virus replication^59^ and mouse hepatitis virus RNA synthesis^60^. Therefore, we postulated that proteins identified in the NE STING co-IP experiment could contribute to IIR signaling and the potential links to IRF3/7 transcription factors suggested a signaling network through which STING might influence IIR from the NE.

### STING NE co-IP partners contribute to IIR

To test whether putative partner proteins identified in the NE STING co-IP experiment are involved in dsDNA triggered IIR, we used a dual-luciferase reporter system in combination with siRNA mediated knockdown of 7 partner proteins with links to IRF3/7 (Fig. 4a) to test for effects on expression of an IFNβ promoter-driven reporter or a reporter activated by NFκB binding. The NFκB- and IFNβ-luciferase reporters are activated upon co-transfection of STING and cyclic GMP-AMP synthase (cGAS) (Fig. 4b). cGAS produces a second messenger (cGAMP) that is bound by STING during IIR^8^ and HEK293FT cells were used because they do not express cGAS (Supplementary Fig. S1e) so that the only source was the transfected plasmid. The cells were also co-transfected with a Renilla luciferase reporter under a thymidine kinase promoter to allow for normalization of transfection efficiency and cell number. Using this assay siRNA knockdown of MEN1, DDX5, snRNP70, and RPS27a all caused a statistically significant drop in IFNβ promoter driven luciferase expression (Fig. 4c), while SYNCRIP, MEN1, DDX5, snRNP70, RPS27a and AATF all exhibited a statistically significant drop in NFκB activated luciferase expression (Fig. 4d). This suggests that these putative STING partner proteins can themselves contribute to IIR. It is interesting that SYNCRIP and AATF were more restricted in only being able to affect luciferase expression from the NFκB driven reporter.

**Fig. 4.**
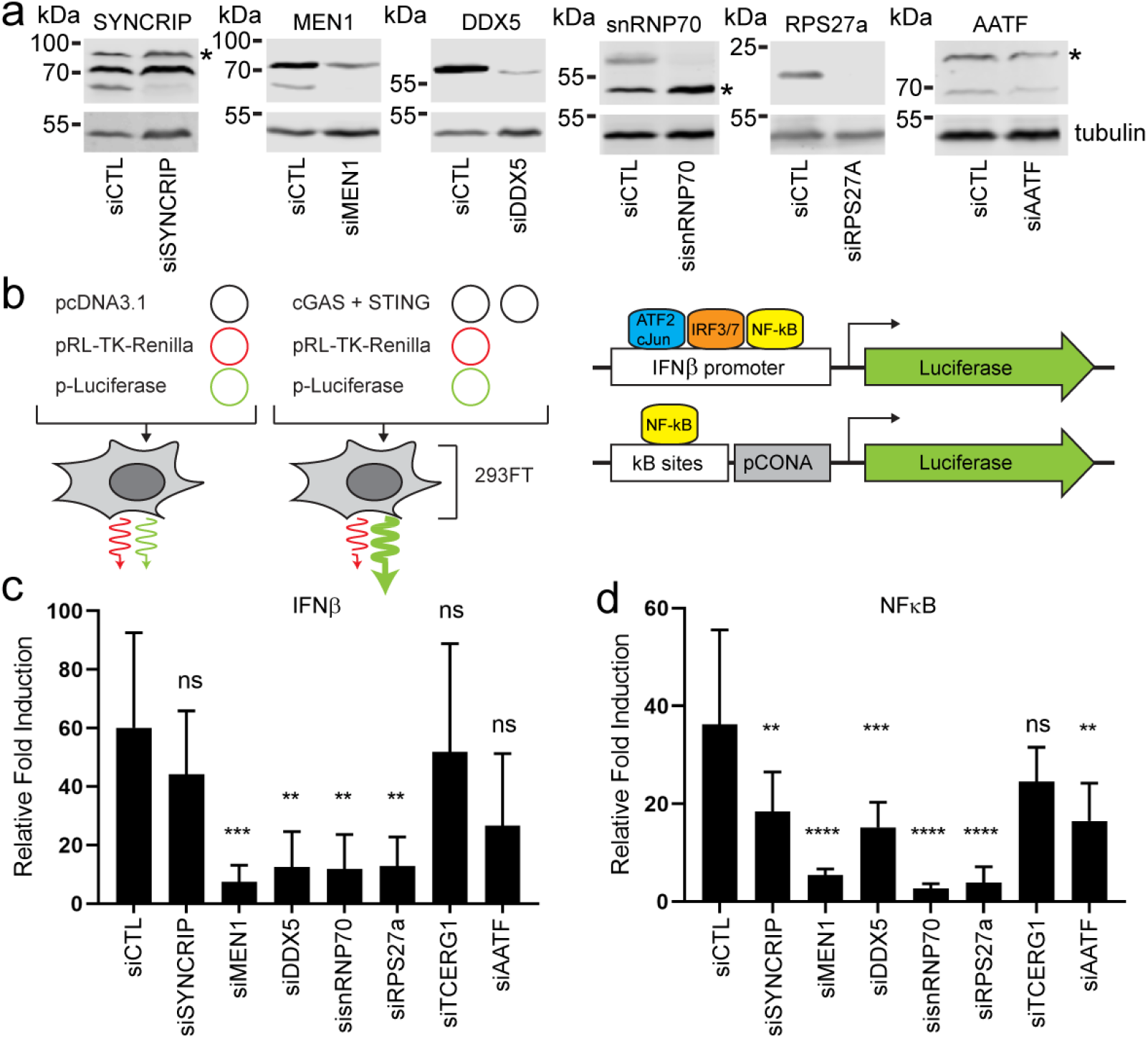
STING putative NE partners contribute to IIR activation. **a** Confirmation of siRNA knockdowns for testing effects of partners in IIR activation assays. Representative Western blots for partners with antibodies that detected proteins of expected molecular weight are shown. * indicates non-specific bands recognised by antibody. In the case of SYNCRIP the highest molecular weight band likely represents the homologous hnRNP R protein which shares a large degree of sequence identity with SYNCRIP and is reported to be recognised by anti-SYNCRIP antibodies. **b** Schematic of dual luciferase assay employed to measure activity of IIR reporter genes. Plasmids expressing Renilla Luciferase variant under a thymidine kinase promoter and Firefly Luciferase under a promoter activated by NFκB binding or the IFNβ promoter are transfected with or without cGAS and STING into 293FT cells. These cells do not express cGAS and have low levels of endogenous STING so the transfection induces IIR in a controlled manner. Comparing the Renilla and Firefly Luciferase levels further controls for differences in transfection efficiency and cell number. **c** IFNβ promoter reporter reveals a significant reduction in the IIR activation when 4 of the 7 STING putative NE partners were knocked down. Six replicates were done with ordinary one-way ANOVA and Dunnett’s multiple comparisons test, ** p ≤ 0.01, *** p ≤ 0.001. **d** NFκB activated reporter reveals a significant reduction in the IIR activation when 6 of the 7 STING putative NE partners were knocked down. Six replicates were done with ordinary one-way ANOVA and Dunnett’s multiple comparisons test, ** p ≤ 0.01, *** p ≤ 0.001, **** p ≤ 0.0001.

To further confirm the role of these STING NE co-IP partners in IIR, independent of the luciferase assay system we measured transcripts of IFNβ with the various knockdowns in HT1080 cells (Supplemental Fig. S3a) +/- initiation of IIR with plasmid DNA or poly(I:C). HT1080 cells have a functional cGAS-STING pathway as shown by the redistribution of STING into perinuclear foci upon dsDNA triggered immune stimulation and the accumulation of IRF3 in the nucleus (Supplementary Fig. S2a and b). These cells also respond to dsRNA triggered immune stimulation as shown by the accumulation of IRF3 in the nucleus following poly(I:C) transfection, indicating that RIG-I/MDA-5/MAVS signaling pathways are functional (Supplementary Fig. S2a)). siRNA knockdown of STING partners SYNCRIP, MEN1, and SNRNP70 did not affect STING or cGAS protein levels as determined by Western blot (Fig. 5a and b). However, knockdown of DDX5 caused a modest reduction in STING protein levels, suggesting that either DDX5 is required for STING stability or an off target effect of the siRNA used. As expected, STING knockdown strongly reduced the amount of IFNβ induction upon plasmid DNA but not poly(I:C) stimulation of IIR (Fig. 5c-e). SYNCRIP, MEN1 and snRNP70 knockdown all reduced IFNβ induction by more than 50% (Fig. 5c and d) an effect that was more marked at 8h post-DNA transfection. Surprisingly, DDX5 knockdown significantly enhanced IFNβ induction with both DNA and poly(I:C) immune stimulation. This effect is especially interesting given that DDX5 siRNA treatment caused a reduction in STING protein levels and suggests that DDX5 indirectly functions as a negative regulator of both DNA and RNA triggered IIR. Surprisingly, despite that several of these STING NE coIP partners are RNA-binding proteins, when IIR was induced with poly(I:C) the other proteins tested did not have a significant effect on IFNβ induction suggesting that they specifically modulate IFNβ induction during DNA triggered immune responses (Fig. 5e).

**Fig. 5.**
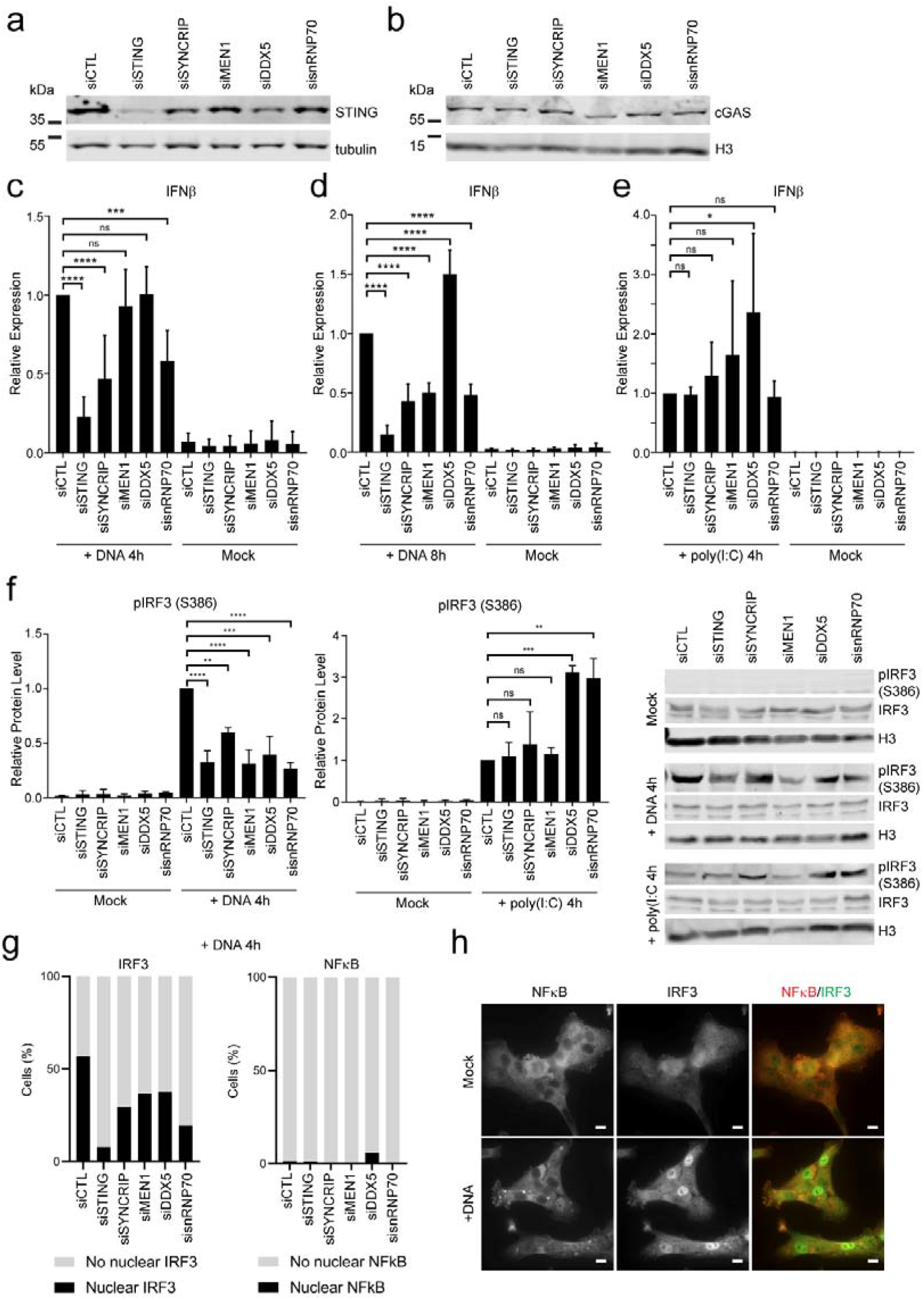
STING putative NE partners have stronger effects on dsDNA-than dsRNA-stimulated IIR. Contributions of STING putative NE partners to IIR were further confirmed by measuring effects on IIR induced by treatment with dsDNA or poly(I:C), dsRNA mimic. These assays were performed in HT1080 cells that express endogenous STING and cGAS. **a** Western blotting confirms minimal effects of siRNAs on STING and (**b**) cGAS expression. Except for siDDX5 which caused a modest reduction in STING protein levels. Representative blots of three independent experiments. **c** Quantification of IFNβ transcripts by qPCR reveals strong effects of STING putative partners, SYNCRIP and SNRNP70 4 h after transfection of dsDNA (n = 3). **d** Effects on IFNβ transcripts are greater 8 h after transfection of dsDNA. SYNCRIP, MEN1, and SNRNP70 siRNAs all reduced IFNβ transcripts while DDX5 siRNA treatment caused a significant increase in IFNβ transcripts relative to siRNA control treated samples (n = 3). **e** In contrast, no effect was observed in response to poly(I:C), except for DDX5 knockdown which caused a significant increase in IFNβ levels (n = 3). (Ordinary oneway ANOVA with Dunnett’s multiple comparisons test * ≤ 0.05, p ** p ≤ 0.01, *** p ≤ 0.001, p **** ≤ 0.0001). **f** Affect of partner protein knockdown on IRF3 phosphorylation (pIRF3) following immune stimulation with dsDNA or poly(I:C). Western blotting representative of three independent experiments. **g** Quantification of the number of cells with accumulation of IRF3 or NNB transcription factors in the nucleus following treatment with siRNAs against STING and putative NE partners and immune stimulation with dsDNA (4 h post-transfection) (n ≥ 100 cells). **h** Representative images for IRF3 and NFκB immunofluorescence used to quantify percentage of cells with nuclear accumulation of IRF3 and NFκB in *g*.

Given the potential links between NE STING partners and IRF3/7 and the effects on IFNβ induction we decided to look at whether their knockdown affected IRF3 activation as measured by IRF3 phosphorylation. IRF3 phosphorylation upon treatment with plasmid DNA was reduced when STING or its NE co-IP partners were knocked down compared to cells treated with control siRNA (Fig. 5f). In contrast, with poly(I:C) treatment there were no obvious effects on IRF3 phosphorylation in cells knocked down for STING and MEN1; however, it was significantly enhanced in cells knocked down for DDX5 and snRNP70, while also slightly increased in cells knocked down for SYNCRIP although not at a statistically significant level (Fig. 5f). As another measure of IRF3 activation, cells treated with siRNAs against STING and partner proteins were assayed for accumulation of IRF3 in the nucleus by microscopy. In agreement with the reduction in phosphorylated IRF3 seen in cells treated with siRNAs against STING and partners following immune stimulation with dsDNA, the percentage of cells positive for accumulation of IRF3 in the nucleus was reduced in all conditions compared to cells treated with a control siRNA (Fig. 5g and Supplementary Fig. S3b). Nuclear accumulation of NFκB (RelA) was also tested, revealing that in response to dsDNA treatment RelA is only weakly activated (phosphorylated RelA accumulates in the nucleus) in HT1080 cells (Fig. 5g and h). In contrast treatment of knockdown cells with poly(I:C) led to a robust activation of IRF3 and NFκB, as determined by the accumulation of NFκB (p65) in the nucleus (Supplementary Fig. S3c).

### STING NE co-IP partner SYNCRIP is antiviral against influenza A virus

Although none of the STING co-IP partners tested were found to have negative effects on poly(I:C) stimulated IFN expression (Fig. 5e), we decided to test whether they might play a role in IIR against an RNA virus, since STING may function in IIR triggered by RNA virus infection in a manner independent of IFN induction^7, 13, 22^. Following siRNA mediated knockdown of STING NE partners, HT1080 cells were infected with the nuclear replicating RNA virus, influenza A virus (IAV). Knockdown of SYNCRIP resulted in significantly higher viral titers as determined by plaque assays, both at low and high multiplicity of infection (MOI) (Fig. 6a and b). This effect was stronger for a mutant IAV (PR8 – N81^61^) which expresses an NS1 protein with a deletion of the effector domain and so is less able to antagonise host IIR (Fig. 6c). To determine whether SYNCRIP expression is altered during viral infection, cell lysates were harvested at multiple time points during infection and blotted for SYNCRIP, revealing no obvious difference in SYNCRIP protein levels during infection (confirmed by presence of viral proteins NP and NS1) compared to mock infected cells (Fig. 6d). STING knockdown has previously been shown to affect IAV replication^18, 62^ and we replicated this here in HT1080 cells. Knockdown of STING resulted in significantly higher viral titers compared to cells treated with a control siRNA (Fig. 6e). Further, we have found that the phenotypic effects of SYNCRIP knockdown replicate those of STING knockdown.

**Fig. 6.**
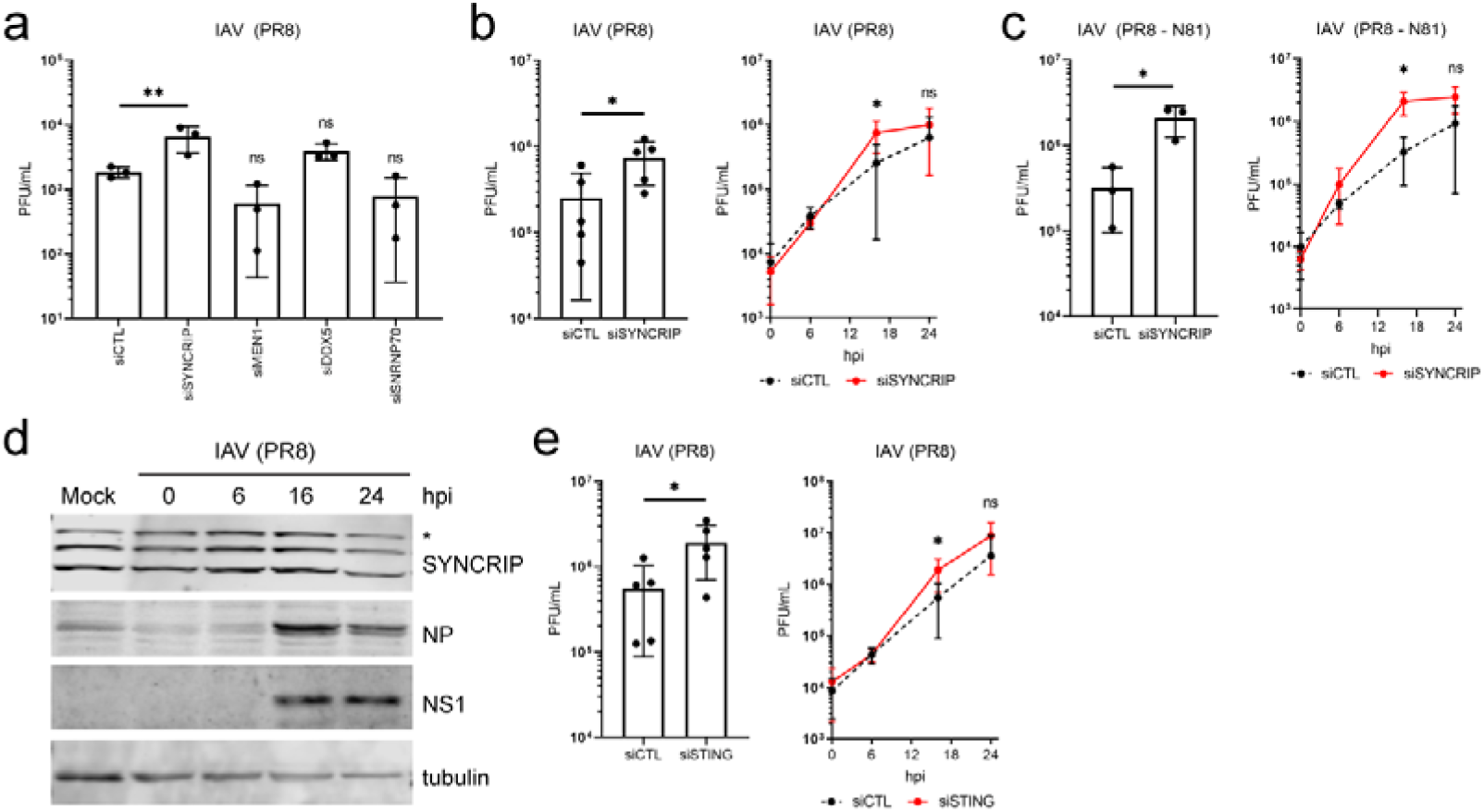
SYNCRIP antagonizes IAV infection. **a** Determination of viral titers in cell culture medium collected from HT1080 cells knocked down for STING partners, 24 hours post infection (hpi) with IAV (PR8 strain) at a multiplicity of infection (MOI) of 0.01, by plaque assay (PFU = plaque forming units) (n = 3) (Ordinary one-way ANOVA with Dunnett’s multiple comparisons test ** p ≤ 0.01). **b** Confirmation of SYNCRIP knockdown effect on IAV titers in cells infected with a higher multiplicity of infection (MOI = 3). Left panel shows significantly higher viral titers at 16 hpi, right panel shows timecourse of infection (n = 5) (student’s T test, * p ≤ 0.05). **c** Effect of SYNCRIP knockdown on viral titers is stronger on IAV mutant virus with truncated NS1 protein (NS1-N81) (MOI = 3). Left panel shows significantly higher viral titers at 16 hpi, right panel shows timecourse of infection (n = 3) (student’s T test, * p ≤ 0.05). **d** Western blotting of SYNCRIP (* indicates non-specific band) and viral proteins, NP and NS1, during IAV infection shows no obvious effect on SYNCRIP protein levels. **e** Confirmation that STING knockdown is beneficial to IAV infection in HT1080 cells as determined by increased viral titers relative to siRNA control treated cells. (MOI = 3). Left panel shows significantly higher viral titers at 16 hpi, right panel shows timecourse of infection (n = 3) (student’s T test, * p ≤ 0.05).

## Discussion

This study identified several proteins that co-IP with the NE pool of STING and found that several of these putative STING partners can themselves contribute to IIR. Specifically, we identified SYNCRIP, MEN1, snRNP70, RPS27a, DDX5, and AATF as novel modulators of IIR. Furthermore, this study directly shows for the first time that part of the endogenous NE pool of STING is present in the INM and that this pool becomes more mobile and redistributes during IIR triggered by both dsDNA and dsRNA stimuli.

In the best characterized STING pathway, recognition of cytoplasmic dsDNA by cGAS triggers cGAMP association with STING to promote its activation. Activated STING dimers translocate from the ER to the Golgi where they accumulate in perinuclear aggregates^6, 7, 8, 63, 64, 65^. From here STING dimers oligomerize, inducing TANK-binding kinase 1 (TBK1) activation which *trans* phosphorylates itself and neighboring STING dimers^15, 66, 67^, leading to the recruitment and activation of IRF3 and eventually the induction of type-I IFNs and pro-inflammatory cytokines. However, STING translocation to or from the nucleus in IIR was previously unknown. Here we have shown for the first time that endogenous STING is clearly present in the INM and that this NE STING pool increases mobility and translocates from the INM to ONM upon IIR activation with the dsRNA mimic poly(I:C) or dsDNA. In light of recent reports that most cGAS is in the nucleus^3, 29, 31, 34^, this raises the possibility that INM STING could be activated before ER STING following detection of nuclear-localized viral dsDNA. Indeed, our finding that STING mobility in the NE increases during infection with the dsDNA nuclear-replicating virus HSV-1 would support this notion. This finding also has implications for a recent report of cGAS-independent activation of STING following detection of DNA damage, in which the authors propose a non-canonical signaling complex composed of STING, TRAF6, IFI16, and p53 that forms in response to DNA damage sensed by PARP1 and ATM, and initiates an NFκB dominated transcriptional response^25^. It is possible that such a signaling complex forms in the nucleus given that the authors of this study reported no redistribution of ER resident STING to perinuclear foci following the induction of DNA damage by etoposide. It is interesting in this regard that one of the more abundant STING NE coIP hits was the DNA damage response protein PARP1 (see Supplementary Table S1). Although this was not included as a top hit because there were only twice as many spectra in the NET23 sample as in the mock when our cutoff was 3-fold, this and other proteins identified in the proteomics further support a role of nuclear STING functioning to sense nuclear DNA damage to induce immune responses in cancer.

Furthermore, we find that the dsRNA mimic poly(I:C) increases STING mobility in the NE and similarly promotes its redistribution from the INM to the ONM. STING protection against RNA viruses seems to function in a different pathway from the well-characterized dsDNA route since STING does not redistribute from the ER to Golgi perinuclear aggregates with poly(I:C) treatment. STING also is reported to not directly bind RNA or poly(I:C)^68^. It was recently reported that STING restricts the replication of RNA viruses through a proposed mechanism dependent on the cytosolic dsRNA sensor RIG-I, and due to a general inhibition of translation independent of PKR and translocon functions^13^. Several of the STING NE partners we identified here could potentially mediate STING effects on translation (e.g. RPS27a, SYNCRIP, snRNP70); however, it also is possible that these partners could provide specific recognition of different RNA viruses and thus serve to provide a variety of novel IIR nucleotide sensors or adaptors. Interestingly, despite having no effect on poly(I:C)-mediated interferon expression, we find SYNCRIP to play a role in antagonizing IAV. Whether this is through a general translation effect, or indirectly through the canonical cGAS-STING pathway stimulated by mitochondrial stress and DNA leakage as reported for other RNA viruses over the years^69, 70^, remains to be determined.

That the INM STING pool can mobilize to translocate to the ONM through the peripheral channels of the NPC may reflect a backup mechanism to signal IIR responses using the peripheral NPC channels when viruses inhibit central channel transport. Viruses often target the central channel of the NPCs to either block transport or to usurp it so that virus transcripts are preferentially transported over host-directed transport^71, 72, 73, 74^, but the peripheral channels are normally used for membrane protein transport^38, 42, 43, 75^ so that STING as a multi-spanning transmembrane protein could bypass this block to signal IIR. This does not preclude the well-established STING signaling cascades from the ER/Golgi compartment normally using the central channel of the NPC — indeed IRF3 is known to translocate through the NPC central channel^76, 77^, but our findings of increased STING mobility and nucleo-cytoplasmic shuttling during IIR through the peripheral NPC channels suggest that STING may provide a backup system for activating IIR when central channel transport is disrupted. Moreover, the 17 STING NE co-immunoprecipitation partners identified here that are known to themselves interact with six IRF3/7 partners could potentially contribute to such IIR activation, enhancing STING functions through a multiply redundant backup system.

The identification of these putative STING partners through the reverse-crosslinking approach does not necessarily mean they directly bind STING and confirming such interactions may require finding particular conditions for each. Moreover, the increasing complexity of STING interactions and its multiple pathways for activating IIR make confirmation of STING’s involvement in their contributions to IIR difficult. Nonetheless, these new putative STING partners clearly can contribute to IIR from the measures shown here; indeed, several reports in the literature further suggest these proteins play roles in IIR to different viruses when re-evalutated in light of our results.

MEN1 binds and represses the activity of the AP1 transcription factor JunD^78^ and the related cJun is an IIR activator^79^, possibly explaining its functioning in IIR. MEN1 was also recently found to affect promoter fidelity at the interferon-gamma inducible IRF1 gene^80^. Interestingly, this function of MEN1 involves its functioning in a complex with the major histone K4 methyltransferase, MLL1, which was also identified as a STING NE co IP partner. STING interaction with this methyltransferase complex could also contribute to the other reported nuclear function for STING in chromatin compaction^28^. In addition, several bromodomain proteins identified as putative STING partners here (e.g. BRD2, BRD3) could also explain this chromatin compaction function or, more excitingly, chromatin remodeling reported to occur in IIR^45^. Furthermore MEN1 is a tumor suppressor^81^ and as such could contribute to reported STING roles in DNA damage sensing in cancer^3, 29, 31, 34, 82, 83^. A recent study showed that MEN1 depletion results in mis-regulation of the p53 pathway leading to increased levels of chromosomal instability and accumulation of DNA damage^83^.

The STING-NE co-IP partners that bind RNA also have several previously reported functions that would be consistent with their ability to support IIR indicated here. For example, DDX5 is targeted by the N(pro) protease of pestivirus, presumably to counter host antiviral defenses^50, 51^. At the same time, DDX5 can be a negative regulator of IFN responses. Our finding that DDX5 knockdown increases type-I IFN expression following DNA or poly(I:C) transfection and IRF3 phosphorylation following poly(I:C) transfection is consistent with a recent report that DDX5 suppressed IFN responses triggered by VSV infection^84^. This study also showed that DDX5 knockdown increased IRF3 phosphorylation, but without testing whether DDX5 knockdown influences IRF3 phosphorylation triggered by a DNA ligand as we do. Our findings here that DDX5 knockdown reduces dsDNA-induced IRF3 activation while elevating IFNβ expression might appear contradictory without the context of these other studies suggesting it can be both pro and anti IIR. Regardless, these results strongly suggest that DDX5 may contribute a regulatory function to IIR. While this study has focused on characterising potential IIR activity of the specific binding partners highlighted for having upstream effects on IRF3/7 transcription factors, it is notable that many more of the top STING NE coIP partners bind RNA and some have previously been shown to mediate IIR. One of these is DDX23 that is a dsRNA sensor recently reported to pair with TRIF or MAVS to mediate IIR^85^.

Other links to viral infection for these newly identified STING partners include the hepatitis B virus HBx protein that alters the intracellular distribution of RPS27a^57^. Also, STING-NE coIP partner AATF is specifically targeted by HIV to impair cellular responses to infection^58^. SYNCRIP/hnRNPQ interestingly facilitates hepatitis virus replication^59^, suggesting an alternate pathway where STING sequestration of this factor might provide another avenue towards host protection from the virus. SYNCRIP was separately reported to interact with the IAV NS1 protein^86^, a major antagonist of the host cell immune response, suggesting the virus may target SYNCRIP due to its positive immune functions. These many RNA-binding partners would provide a highly redundant backup system so that knockout of any single one would only moderately impact on IIR, consistent with the moderate but significant reduction in IIR signaling observed for SYNCRIP knockdown. Our data are consistent with the notion that STING plays a wider role in signaling than its initial description as an adaptor in cytosolic DNA sensing, initiating different responses based on diverse inputs from DNA damage^25^ to RNA virus infection^13^. The wide range of STING nuclear partners combined with its ability to translocate out of the nucleus with treatments that activate IIR potentially provides a valuable redundancy and novel mechanism for STING functions that could better elucidate how it protects cells against both RNA and DNA viruses.

## Methods

### Cell culture and transfections

HT1080, MDCK, HEK293T and HEK293FT cells were maintained in high glucose DMEM (Lonza) supplemented with 10% fetal bovine serum (FBS), 100 μg/μl penicillin and 100 μg/μl streptomycin sulfate (Invitrogen). To generate a stable inducible NET23/STING expressing cell line, a lentivirus vector encoding doxycycline inducible NET23/ STING fused to GFP at the C-terminus (pLVX-TRE3G backbone, Clontech) was prepared by standard procedures and transduced into HT1080 cells. Transduced cells were selected with geneticin at 500 μg/ml. Inducible stable cells were treated with 0.05 μg/ml doxycycline for 20 h in order to express NET23/STING expression before use in experiments.

### IIR induction

To induce IIR in 6 well-plate format for immunoblotting and qPCR experiments, cells were transfected with 10 μg poly(I:C) (Sigma), 5 μg pcDNA3.1, or transfection reagent alone (mock) using 4 μl Lipofectamine 2000 transfection reagent in 200 μl optiMEM added to cells in 2 ml culture medium. For immunofluorescence staining to quantify nuclear IRF3/NFκB, cells were transfected with 2 μg poly(I:C) (Sigma), 1 μg pcDNA3.1, or transfection reagent alone (mock) using 1 μl Lipofectamine 2000 transfection reagent in 50 μl optiMEM added to cells in 0.5 ml culture medium.

### Immunofluorescence microscopy

HT1080 cells seeded on glass coverslips were fixed in 3.7 % formaldehyde for 15 min. Cells were then washed in PBS and permeabilized with 0.2% Triton X-100 for 7 min before blocking in 4 % BSA for 30 min at room temperature. Cells were incubated with primary antibodies in blocking buffer overnight at 4 °C, antibodies used were: anti-STING (AF6516, R&D Systems), anti-Nup153 (QE5, ab24700, Abcam), anti-IRF3 (FL-425, sc-9082, Santa Cruz Biotechnology), anti-NFκB (L8F6, 6956S, Cell Signaling Technology). The following day coverslips were washed with PBS and incubated for 1 h with appropriate secondary antibodies conjugated to Alexa Fluor dyes (MolecularProbes). DNA was visualized with DAPI (4,6-diamidino-2 phenylindole, dihydrochloride) and coverslips mounted in fluoromount G (EM Sciences). For anti-STING (D2P2F, #13647, Cell Signaling Technology) antibody staining, cells were fixed for 15 min in ice cold methanol at −20 °C. Immunofluorescent staining was then carried out as described in^87^. Images were obtained using a Zeiss AxioImager equipped with 1.45 NA 100x objective.

### Structured illumination microscopy

HEK293T cells transfected with STING-GFP were fixed for 7 min in 3.7% formaldehyde, washed with PBS, and then permeabilised 6 min in 0.2% Triton X-100. Cells were then blocked with 10% FBS, 200 mM glycine in PBS, and reacted for 40 min at RT with antibodies to Nup153 (Covance) or Nup352 (kind gift of F. Melchior). All secondary antibodies were Alexa fluor highly immunoadsorbed goat IgG (MolecularProbes). DNA was visualized with DAPI (4,6-diamidino-2 phenylindole, dihydrochloride) and coverslips mounted in Vectashield.

Structured illumination images (Figure 1) were taken on the OMX system at the University of Dundee microscopy facility (details described at http://microscopy.lifesci.dundee.ac.uk/omx/). OMX was employed to determine whether a protein is in the outer or the inner nuclear membrane by co-staining with antibodies to nuclear pore complex (NPC) cytoplasmic filament protein Nup358 and nuclear basket protein Nup153 as these structures extend well beyond the 50 nm spacing between the inner and outer nuclear membranes (INM and ONM respectively).

### Immunogold electron microscopy

HT1080 control cells or HT1080 cells stably transfected with inducible STING-GFP that was induced O/N with doxycycline were fixed for 2 min by the addition of a 2X fixation buffer (8% formaldehyde, 0.2M PHEM buffer) to culture medium, this was then replaced with 1X fixation buffer (4% formaldehyde, 0.1M PHEM buffer, 0.05% glutaraldehyde) and fixation continued for 2h, cells were then scraped and transferred to microfuge tubes, before pelleting and washing with PBS. Cell pellets were then stored in PBS prior to sectioning and immunostaining. The cell pellets were embedded in 10 % gelatine before being infused overnight with 15% PVP and 1.7M sucrose in 0.1M phosphate buffer. Liquid nitrogen frozen pellets were sectioned on a cryo-ultramicrotome (UC7 with FC7 cryo-attachment; Leica). Immunoelectron microscopy was performed using the Tokuyasu method^88^. Cryosections were thawed, rinsed with 1 % glycine in PBS with and blocked with 1% BSA in PBS. For endogenous STING, grids were incubated with sheep anti-STING antibody at 1:7 dilution (AF6516, R&D Systems), rinsed in PBS, then incubated with a 6nm donkey anti–sheep IgG antibody conjugated to 6 nm colloidal gold (Aurion). For STING-GFP expressing cells grids were incubated with a rabbit anti GFP antibody at 1:20 dilution (Abcam), rinsed in PBS, then incubated with a 5nm goat anti-rabbit IgG antibody conjugated to 5 nm colloidal gold (Aurion) Grids were then rinsed in PBS, transferred to 1% glutaraldehyde (Agar Scientific) in PBS, washed in water, and embedded in 2% methyl cellulose containing 0.4% uranyl acetate (Agar Scientific). Imaging was performed at 100 kV with a Hitachi H7600 TEM and Xarosa 20 Megapixel camera.

### FRET-FLIM

HEK293T cells were seeded at 70,000 cells on glass coverslips No. 1.5 per well of a 24-well plate. The next day cells were transiently transfected with plasmids encoding Lamin A-GFP, an RFP-GFP tandem construct, or Lamin A-GFP and STING-RFP^27^. After 24 h cells were fixed 15 minutes in 3.7% formaldehyde before washing in PBS and mounting in Vectashield®. Imaging was performed on a Leica SP5 SMD confocal laser scanning microscope equipped with PicoHarp 300 (TCSPC module and picosecond event timer) and single photon avalanche detectors. Leica application suite with FLIM wizard software and integrated Symphotime software were used for single photon counting acquisition and FLIM measurements carried out for 5 min per field of view. FLIM data was analysed using FLIMfit 5.1.1 software (the FLIMfit software tool developed at Imperial College London).

### FRAP

To induce IIR for FRAP experiments, an HT1080 cell line stably transfected with a doxycycline-inducible STING construct was first treated 20 h prior to IIR induction with 0.05 μg/ml doxycycline. For stimulation of IIR cells were plated onto a 6-well plate (200,000 cells/well) and infected with a multiplicity of infection (MOI) of 10 using HSV-1 strain 17+. The virus was added to cells in 0.5 ml culture medium. The cells were incubated for 1 h at 37°C and 5% CO_2_ to facilitate adsorption of the virus. Subsequently 1.5 ml medium was added and the cells were incubated for additional 2 h before FRAP imaging. For poly(I:C) stimulation cells were transfected in 6-well plates with 10 μg polyI:C using Lipofectamine 2000 (Invitrogen) for 2 h before imaging. All fluorescence recovery after photobleaching (FRAP) experiments for GFP-fusion constructs were performed on a Leica SP5 microscope equipped with an Argon laser using the 488 nm laser line and a 60x HCX PLAPO NA 1.4 oil objective. Temperature was maintained at 37°C in an environmental chamber (Life Imaging Services, Switzerland), cells were gassed using 5% CO_2_ in air using a gas mixer (Life Imaging Services). Stably transfected inducible STING-GFP or NET55-GFP HT1080 cells grown on 25 mm round coverslips mounted in an Attofluor incubation chamber (Life Technologies) were first induced for expression of the fusion protein overnight with doxycycline. The next day the coverslips were clamped into the chamber with 2 ml of preheated (37°C), phenol-free complete DMEM containing 25 mM Hepes-KOH. Five prebleach images were taken followed by bleaching a spot of 1 μm for 1 s at full laser intensity. Subsequent images were taken in 2 phases. The first rapid phase consisted of 25 frames every 0.65secs to observe the rapid initial recovery. The second slow phase consisted of 75 frames every 2 s to observe the slower final stages of recovery, these parameters were shown by preliminary experiments (data not shown) to allow complete steady state recovery in all cells.

Image data was processed using Image-Pro Premier (Media Cybernetics Inc., MD, USA). Background and photobleach corrections were engaged using an algorithm written by D.A.K. according to^89^. A macro was written in VB.Net within Image Pro Premier whereby a region of interest (ROI) was applied to the bleach spot, background and non-bleached area of a nearby cell and corrected for movement automatically compared to the 5 pre-bleach images. The t½s were calculated from the normalized fluorescence values.

### Single-Molecule Fluorescence Recovery After Photobleaching (smFRAP) microscopy

Imaging was performed using an Olympus IX81 equipped with a 1.4-NA 100x oil-immersion apochromatic objective (UPLSAPO 100XO, Olympus, Center Valley, PA). An Obis™ solid state 488-nm (Coherent Inc, Santa Clara, Ca) was passed through dichroic filters (Di01-R405/488/561/635-25×36, Semrock, Rochester, NY) and emission filters (NF01-405/488/561/635-25×5.0, Semrock, Rochester, NY) as well a circular variable metallic neutral density filter (Newport, Irvine, CA) and directed into the microscope using a micrometer stage (Newport, Irving, CA). Cells modified to express doxycycline inducible STING-GFP were plated on No. 0 cover glass 35 mm petri dishes with 14 mm Microwell (MatTek, Ashland, MA). Cells were transfected via lipofection agent Transit-X2 (Mirus Bio LLC, Madison, WI) 2 h prior to imaging for plasmid DNA or poly(I:C), or transfection reagent alone (mock). Growth media was replaced with Transport Buffer (20 mM HEPES, 110 mM KOAc, 5 mM NaOAc, 2 mM MgOAc, and 1 mM EGTA, pH 7.3) 30 min prior to imaging to slow membrane movements and reduce autofluorescence. Imaging was performed on the Nuclear Envelope region opposite the Endoplasmic Reticulum. The region was photobleached for 1 minute at 5 mW and then imaged for 30 seconds at 50 μW. An optical chopper rotating at 2-Hz was utilized during capture to allow for fluorescence recovery. Images were captured using the Slidebook software package (Intelligent Imaging Innovation, Denver, Co). Data were analyzed using the FIJI ImageJ^90^ plugin GDSC SMLM (Single Molecule Light Microscopy ImageJ Plugins, University of Sussex, http://www.sussex.ac.uk/gdsc/intranet/microscopy/UserSupport/AnalysisProtocol/imagej/gds_c_plugins/, 2013; Herbert, A. *Single Molecule Light Microscopy ImageJ Plugins*, 2014) and OriginPro 2019 (OriginLab, Northampton, MA).

### Nuclear-specific cross-linking co-IP

2 x 10^8^ HEK293T cells were transfected using Ca2PO4 with either STING-GFP or mock transfected. As overexpression of STING tends to eventually result in apoptosis, cells were taken at ~20 h post-transfection. Note that a mock-transfected cell population was prepared for reversible crosslink immunoprecipitations the same way as the transfected cells. Cells were first rinsed on the plates with PBS, then incubated with trypsin (concentration) for 3 min and shaken from the plates. Recovered cells were washed in 10% FBS in PBS to inactivate trypsin and pelleted by centrifugation at 250 x g for 10 min at room temperature. The NE isolation followed published protocols^36, 91^. For all subsequent steps the following protease inhibitors were added freshly to solutions: 1 mM AEBSF [4-(2-Aminoethyl) benzenesulfonyl fluoride hydrochloride], 1 μg/ml aprotinin, 1 μM pepstatin A, and 10 μM leupeptin hemisulfate. Cells were then washed with PBS, repelleted and next resuspended in roughly 100x the pellet volume with hypotonic lysis buffer (10 mM HEPES pH 7.4, 1.5 mM MgCl_2_, 10 mM KCl with freshly added 2 mM DTT and protease inhibitors). Cells were allowed to swell on ice for 5 min and then nuclei were released by dounce homogenization with 10 vigorous strokes using a type B pestle (Wheaton, clearance between 0.1 and 0.15 mm). Thereupon 1/10 volume of 1 M KCl and 1/10 volume of 2.2 M sucrose were immediately added. Nuclei were pelleted through a 0.9 M sucrose cushion in the same buffer with the salt (0.9 M sucrose, 10 mM HEPES pH 7.4, 1.5 mM MgCl_2_, 110 mM KCl with freshly added DTT) at 2,000 x g in a swinging bucket rotor (e.g. 4,000 rpm in a Beckman Coulter J6-MC floor model centrifuge) for 20 min at 4°C. Nuclear pellets were resuspended at 1.5 million nuclei/ ml in 0.25 M sucrose, 10 mM HEPES pH7.4, 10 mM KCl, 2 mM MgCl2, 1.5 mM CaCl_2_, with 2 mM DTT, protease inhibitors and 4 U/ml DNase and 1 μg/ml RNase and incubated at room temperature to digest chromatin while swelling nuclei to release the digested chromatin. These were then pelleted through a 0.9 M sucrose cushion as before only at 6,000 x g using a swinging bucket rotor for 20 min at 4°C. The process was repeated with increasing the DNase to 20 U/ml and the RNase to 10 μg/ml and pelleted as above to generate a crude NE fraction, though this time the pellet was resuspended in the same buffer lacking the DNase, RNase, and DTT.

During the DNase/RNase incubations the orthophenanthroline copper was prepared by mixing a solution of 200 mM CuSO_4_ in water at 1:1 with 400 mM 1,10-orthophenanthroline in ethanol. The crosslinker complex is allowed to form by gentle rotation for 30 min at room temperature. The orthophenanthroline copper solution was then added at 1:100 to the NEs resuspended without DTT and incubated for 30 min at 30°C. The reaction was stopped by addition of EDTA to 5 mM and the crosslinked NEs were pelleted at 6,000 x g using a swinging bucket rotor for 20 min at 4°C.

The pellet was resuspended in RIPA buffer with freshly added protease inhibitors and incubated for 20 min at room temperature. This material was then sonicated on ice for 5 min using a bath sonicator with a rotating cycle of 15 s on and 15 s off. Large insoluble material was removed by pelleting at 100 x g for 1 min and antibodies were added for immunoprecipitations. Both mock transfected lysate and STING-GFP transfected lysate were separately incubated with antibodies against GFP (Life Technologies A11122). All antibodies were incubated overnight at 4°C with gentle rotation. The next day antibody-complex conjugates were incubated with Protein A-sepharose beads and washed on the beads with 0.5x RIPA buffer with freshly added protease inhibitors 3 times. Complexed proteins were then released from the beads with 50 mM DTT incubated at room temperature for 30 min and material processed for mass spectrometry.

### SDS-PAGE analysis and in-gel digestion for mass spectrometry

Released proteins were in-gel digested as described elsewhere^92^. In brief, a band of coomassie-stained gel was excised and the proteins where digested using trypsin and proteins were reduced in 10 mM DTT for 30 min at 37°C, alkylated in 55 mM iodoacetamide for 20 min at room temperature in the dark, and digested overnight at 37°C with 12.5 ng/μL trypsin (Proteomics Grade, Sigma). The digestion media was then acidified to 0.1% of TFA and spun onto StageTips as described in the literature^93^. Peptides were eluted in 20 μL of 80% acetonitrile in 0.1% TFA and were concentrated to 4 μL (Concentrator 5301, Eppendorf AG). The peptides sample was then diluted to 5 μL by 0.1% TFA for LC-MS/MS analysis.

### Mass spectrometry analysis

An LTQ-Orbitrap mass spectrometer (Thermofisher Scientific) was coupled on-line to an Agilent 1100 binary nanopump and an HTC PAL autosampler (CTC). The peptides were separated using an analytical column with a self-assembled particle frit^94^ and C_18_ material (ReproSil-Pur C18-AQ 3 μm; Dr. Maisch, GmbH) was packed into a spray emitter (100-μm ID, 8-μm opening, 80-mm length; New Objective) using an air-pressure pump (Proxeon Biosystems). Mobile phase A consisted of water, 5% acetonitrile, and 0.5% acetic acid; mobile phase B, consisted of acetonitrile and 0.5% acetic acid. The gradient used was 98 min. The peptides were loaded onto the column at a flow rate of 0.7uL/min and eluted at a flow rate of 0.3uL/min according to the gradient. 0% to 5%B in 5 min, 5% to 20% buffer B in 80 min and then to 80% B in 13 min. FTMS spectra were recorded at 30,000 resolution and the six most intense peaks of the MS scan were selected in the ion trap for MS2, (normal scan, wideband activation, filling 7.5E5 ions for MS scan, 1.5E4 ions for MS2, maximum fill time 150 msec, dynamic exclusion for 150s sec). Searches were conducted using Mascot software (Version 2.2.0) against a database containing Human sequences (sprothuman201106). The search parameters were: MS accuracy, 6 ppm; MS/MS accuracy, 0.6 Da; enzyme, trypsin; allowed number of missed cleavages, 2; fixed modification, carbamidomethylation on Cysteine; variable modification, oxidation on Methionine.

### siRNA knockdowns

For HT1080 and HEK293FT cells, one day prior to knockdown cells were seeded at 300,000 cells/well of a 6-well plate. The next day siRNA knockdowns were performed using siRNA duplexes listed in table 2 and jetPRIME transfection reagent (POLYPLUS) according to the manufacturer’s instructions for a final concentration of 50 nM siRNA, 4 μl jetPRIME, 200 μl buffer per well. For STING knockdown a mix of siRNA#1 and #2 were used for a final concentration of 50 nM. 24 h later cells were trypsinised and reseeded 1:2 in 6 well plates for western blotting and qPCR experiments, or at 50,000 cells/well of a 24-well plate on 13mm No. 1.5 glass coverslips for immunofluorescence.

**Table 2 –.**
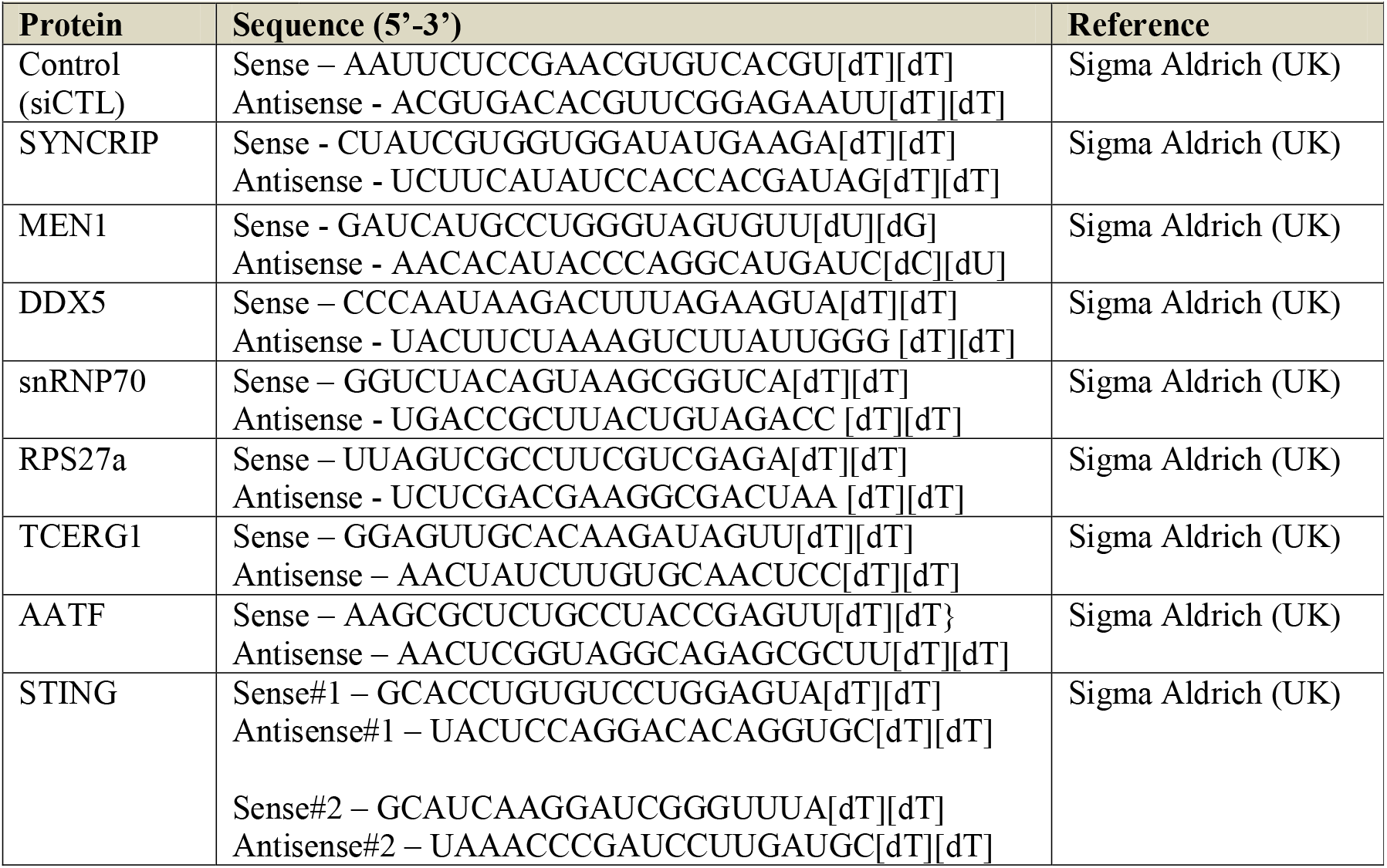
siRNA duplexes

### Dual-luciferase reporter assay

Following siRNA knockdown transfection in 6-well plates as described for HT1080 cells, 5×10^4^ cells were seeded per well of a 48-well plate. After 24h, cells were co-transfected with plasmids expressing luciferase reporter constructs (IFNβ-firefly or NF-κB-firefly at 30ng and Prl-TK-renilla at 5ng) and either cGAS and STING (20ng each) or empty vector DNA (pcDNA3.1 at 40ng). For each well, 1 μL of Fugene (Promega) was incubated with 25 μL of Opti-MEM for 5 min at RT before the addition of plasmid constructs and 280 ng of the required siRNA oligos (second knockdown). This final mixture was incubated for a further 20 min at RT before addition to each well containing 150 μL fresh media. After 24 h media was exchanged for 300 μL fresh media.

To measure luminescence produced by luciferase activity, cells were harvested 96 h after the initial siRNA knockdown transfection. Media was removed and cells were washed in PBS before re-suspension in 75 μL 1X Passive Lysis Buffer (PLB; Dual-Luciferase Reporter Kit, Promega). Cells were incubated in PLB for 15 min at RT and further homogenised by pipetting. 25 μL of the homogenised lysate was transferred to a well on a 96-well plate and luminescence signal detected using a Modulus Microplate Multimode Reader (Turner Biosystems). The plate reader was programmed to inject 40 μl of Luciferase Assay Substrate re-suspended in Luciferase Assay Buffer II and 35 μl of 1X Stop&Glo Reagent (Dual-Luciferase Reporter Kit, Promega) per well, with a 2s delay between injections and 10 s measurement period after each injection. Luminescence signal produced by IFNβ/NF-κB-firefly luciferase was divided by pRL-TK-renilla luciferase luminescence signal to control for variation in transfection efficiency and cell number.

### Bioinformatics analysis

Gene ontology (GO) analysis was performed in R using the Bioconductor package GOstats^95^. Significantly over-represented categories and other biologically interesting classes were represented as a piechart. Only experimentally verified GO-functional annotations were used, including EXP (Inferred from EXPeriment), IDA (Inferred from Direct Assay), and IPI (Inferred from Physical Interaction). We used the following GO terms: “defense response” (GO:0006952), “immune response” (GO:0006955), “response to virus” (GO:0009615), “apoptosis” (GO:0006915), “cell cycle” (GO:0007049), “regulation of cell cycle” (GO:0051726), “negative regulation of cell proliferation” (GO:0008285), “positive regulation of cell proliferation” (GO:0008284), “regulation of transcription” (GO:0006355), “transcription factor” (GO:0001071), “RNA splicing” (GO:0008380), “RNA binding” (GO:0003723), “DNA binding” (GO:0003677), “chromatin” (GO:0000785), “chromatin binding” (GO:0003682), “chromatin modification” (GO:0016568), “chromatin organization” (GO:0051276). To plot MS data piecharts, the relative proportions of the various classes were calculated from the relative abundances of the genes identified from each GO category, and this was compared to a piechart compiled from all the genes in the human genome considering the same categories. The heatmaps simply highlight which genes have been annotated with each GO term shown.

To map further interactions of proteins co-immunoprecipitated with STING, the proteins were searched against the human protein reference database (HPRD, Johns Hopkins University) to identify primary and secondary interactors, using Cytoscape^96^ for visualisation.

### qPCR

RNA extraction was achieved using RNeasy Mini Kit (Qiagen) as per the manufacturer’s instructions. The quality and concentration of the RNA was then assessed using a NanoDrop 2000c spectrophotometer and run on a TAE gel to check for degradation. If not immediately used, RNA was stored at −80°C in separate aliquots to reduce freeze thawing. RNA was converted into cDNA using SuperScript II reverse transcription reagent (Invitrogen) and quantitative real time-PCR (qRT-PCR) performed with normalization to GAPDH. The cycle program consisted of an initial denaturing step of 5 min at 95°C, then 45 cycles of 10 sec at 95°C, 15 sec at 51°C, and 20 sec at 72°C, followed by a melting curve analysis step. qPCR primers were: IFNβ Fw5’-CCTGAAGGCCAAGGAGTACA-3’ and Rev 5’-AGCAATTGTCCAGTCCCAGA-3’ and GAPDH Fw 5’-GTGAAGGTCGGAGTCAACG-3’ and Rev 5’ATGACAAGCTTCCCGTTCTC-3’.

### Immunoblotting

For immunoblotting cell lysates were collected directly by addition of 2x sample buffer (with the addition of Na_3_VO_4_ and NaF for phosphorylated proteins) to washed cell monolayers. Samples were boiled for 5 min at 95 °C and sonicated 1 sec on 1 sec off with a probe sonicator before separation of proteins by SDS-PAGE. Proteins were transferred to nitrocellulose membranes, blocked with 5% BSA or Milk for 1 h at room temperature before probing with primary antibodies overnight. Primary antibodies were: anti-STING (AF6516, R&D Systems), anti-STING (D2P2F, #13647, Cell Signaling Technology) anti-IRF3 (FL-425, sc-9082, Santa Cruz Biotechnology), anti-cGAS (HPA031700, Atlas Antibodies), anti-cGAS (D1D3G, #15102, Cell Signaling Technology), anti-phospho-IRF3 (S386) (ab76493, Abcam), anti-H3 (ab10799, Abcam), anti-tubulin (GTU-88, T6557, Merck), anti-tubulin (ATN02, Cytoskeleton Inc.), anti-SYNCRIP (7A11.2, MAB11004, Millipore), anti-MEN1 (ab2605, Abcam), anti-DDX5 (ab10261, Abcam), anti-snRNP70 (ab51266, Abcam), anti-RPS27a (ab172293, Abcam), anti-AATF (ab39631, Abcam), anti-Lamin A/C (3262, generated by ECS), anti-NS1 (NS1-RBD, generated by PD), anti-NP (A2915, generated by PD).

### Generation of IAV, virus titrations, and infections

IAV strain A/Puerto Rico/8/34 (PR8) stocks were generated using reverse genetics as previously described^61, 97^. 8 pDUAL plasmids (250 ng each with 4 μl Lipofectamine 2000 (Life Technologies, Loughborough, UK) containing sequences of a complete IAV genome were used to transfect HEK293T cells simultaneously, with transfections either incorporating the WT segment 8 sequence or an NS1 mutant (NS1-N81^61^). After overnight incubation, virus growth medium (DMEM supplemented with 5 □ μg/ml TPCK-treated trypsin, 0.14% BSA fraction V and penicillin (100□U/ml) and streptomycin (100□μg/ml)) was added to allow a small-scale amplification of the viruses in HEK293T cells. After 4Mh, the virus particle-containing supernatants were passaged on MDCK cells to further amplify the viruses to obtain working stocks.

Virus titres were determined by plaque titration on MDCK cells in 6 well plates. Cells were inoculated with virus dilutions for 1 □h, then an overlay (mixture of equal volume of DMEM and 2.4% Avicel (Sigma-Aldrich, UK) supplemented with 1 μg/ml TPCK-treated trypsin and 0.14% BSA fraction V) was put onto the wells. After 4Mh, cells were fixed using 3.7% formaldehyde and stained with 0.01% Toluidine Blue. Virus titres were calculated by plaque count*dilution factor/(volume of inoculum) and expressed as PFU/ml.

HT1080 cells were mock infected or infected with either wild-type PR8 or NS1-N81 mutant virus at an MOI of 0.01 or 3 as indicated in figure legends. Culture media of infected cells was collected at time points given post-infection for use in plaque assays, or cells were harvested in laemmli buffer for Western blotting.

### Reporting summary

Further information on research design is available in the Nature Research Reporting Summary linked to this article.

## Data availability

All mass spectrometry data will be deposited to ProteomeXchange Consortium and MassIVE databases upon acceptance of the manuscript.

## Acknowledgements

We thank Ravi Badwe for assistance in nuclear envelope preparations from transfected mammalian cells, Ines Alvarez-Rodrigo for assistance in optimizing the luciferase reporter assay, Emma Winchester for support at the Dundee OMX facility, and Jessica Valli for assistance with the FRET-FLIM microscope at the Edinburgh Super-Resolution Imaging Consortium (ESRIC) facility at Heriot-Watt University. Funding for this work was principally provided by Wellcome Senior Fellowship 095209 to ECS and grant 092076 for the Centre for Cell Biology and MRC grant MR/R018073 to ECS. Project work by MT and WY was supported by grants from the US National Institutes of Health (NIH GM094041, GM097037, GM116204 and GM22552) to W.Y. Work by ACR and MWG was funded by Biotechnology and Biological Sciences Research Council grant BB/R014094/1. Work by EG and PD was funded through BBSRC Institute Strategic Programme grant BB/P013740/1 and project grant BB/S00114X/1. JR and FLA were funded through Wellcome Trust Senior Fellowship 103139 and equipment grant 091020 to JR. PM was funded by a Royal Society Research Fellowship Award DH051766 and a Royal Society research grant RG090330. CRD was funded through a Wellcome PhD studentship 109089. NSR was funded by a Principal’s Scholarship from the University of Edinburgh. GJT was funded through Wellcome Trust Senior Fellowship 090940 to GJT. EG was supported by Wellcome Trust/ Royal Society Sir Henry Dale Fellowship 211222/Z/18/Z. Finally, the Dundee OMX is supported by MRC Next Generation Optical Microscopy Award MR/K015869/1 and SULSA.

## Author Contributions

Conceived/ designed experiments: CRD, JIH, PM, ECS, GJT, EG, PD. Performed experiments: CRD, PM, NSR, JIH, FLA, MT, ACR, ECS. Provided reagents/ analysis tools: DAK, JR, GJT, MWG, WY, EG, PD. Wrote the manuscript: ECS, CRD.

## Additional information

Supplementary information accompanies this paper including Supplemental Table S1 with detailed listings of all co-immunoprecipitation proteins from reverse crosslinked nuclear envelopes is available in the online version of the paper. Mass spectrometry data have been deposited in ProteomeXchange and MassIVE. The authors declare no competing financial interests. Correspondence and requests for materials should be addressed to ECS

## Author Information

Author contact information:

Charles R. Dixon

Poonam Malik

Jose I. de las Heras

Natalia Saiz Ros

Flavia de Lima Alves

Mark Tingey

Eleanor Gaunt

A. Christine Richardson

David A. Kelly

Martin W. Goldberg

Greg Towers

Weidong Yang

Juri Rappsilber

Paul Digard

Eric C. Schirmer

Street addresses:

JIH, PM, NSR, IAR, DAK, FLA, JR, ECS — Wellcome Trust Centre for Cell Biology, University of Edinburgh, Kings Buildings, Swann 5.22, Max Born Crescent, Edinburgh EH9 3BF, UK

GJT - University College London, Division of Infection and Immunity, 90 Gower Street, London, WC1E 6BT, UK

EG and PD — Division of Infection and Immunity, Roslin Institute, University of Edinburgh, Edinburgh, EH25 9RG, UK

MT and WY — Temple University, Department of Biology, Philadelphia, PA 19122, USA

ACR and MWG — Durham University, School of Biological and Biomedical Sciences, Microscopy and Bioimaging, Mountjoy Science Laboratories, South Road, Durham DH1 3LE, UK

