## Supplemental Figures for "STING Nuclear Partners Contribute to Innate Immune Signalling Responses"

Supplementary Figures

**Supplementary Fig. S1** STING targets to the ER and NE **a** Immunogold EM minus anti-STING antibody staining shows no gold particles, indicating specific labelling of primary anti-STING antibody by secondary antibody conjugated to gold particles. Immunogold EM staining performed against endogenous STING shows STING localisation at the ER as well as nuclear envelope. Immunogold EM staining against stably expressed STING-GFP fusion protein shows that STING-GFP exhibits a similar localisation to endogenous STING, present at both the ER and nuclear envelope. Scale = 500 nm. **b** Confocal microscopy of STING-RFP and Lamin A-GFP constructs used for FRET-FLIM experiment confirm both proteins localise as expected (Lamin A at the nuclear envelope and STING-RFP at the ER and nuclear envelope). Scale = 10 μm.

**Supplementary Fig. S2** Validation of canonical STING function in HT1080 cells and HT1080 cells stably transfected with inducible STING-GFP. **a** Immunofluorescence staining using the same STING antibody (AF6516) as for immunogold EM (Fig. 1 and S1) against endogenous STING in HT1080 cells shows that STING is distributed throughout the ER as well as a pool at the nuclear envelope, highlighted by Nup153 staining, in mock stimulated HT1080 cells. 2 h post-immune stimulation with dsDNA STING accumulates in perinuclear foci and IRF localises to the nucleus, while 2 h post-immune stimulation with poly(I:C) STING is still localised throughout the ER and at the NE, and IRF3 has accumulated in the nucleus as expected. **b** Confirmation of a pool of endogenous STING at the nuclear envelope using a different anti-STING antibody (D2P2F) requiring methanol fixation. **c** STING-GFP expressed in stably transfected HT1080 cells behaves similarly to endogenous STING, distributed throughout the ER and at the nuclear envelope in mock stimulated cells and cells stimulated with poly(I:C), while accumulating in perinuclear foci 2 h post-stimulation with dsDNA. Scale = 10 μm. **d** Verification of anti-STING antibody specificity, both anti-STING antibodies recognise a single protein band by immunoblot that is decreased in abundance after treatment with siRNAs against STING. **e** Verification of intact canonical DNA sensing cGAS-STING pathway in HT1080 cells which express detectable levels of both STING and cGAS compared to HEK293T and HEK293FT cells used in this study which express very low levels of STING and undetectable levels of cGAS.

**Supplementary Fig. S3** STING and NE partner knockdown in HT1080 cells **a** Validation of siRNA mediated knockdown of STING and nuclear envelope co IP partners in HT1080 cells. Immunoblotting representative of three independent knockdowns. * indicates non-specific protein band recognised by antibody. **b** Example images of IRF3 accumulation in the nucleus of cells 4 h post-stimulation with dsDNA, quantified in Fig. 5g. **c** Poly(I:C) stimulation induces the accumulation of both NFκB (p65) and IRF3 in the nucleus of HT1080 cells. Scale = 10 μm.

Supp Fig S1

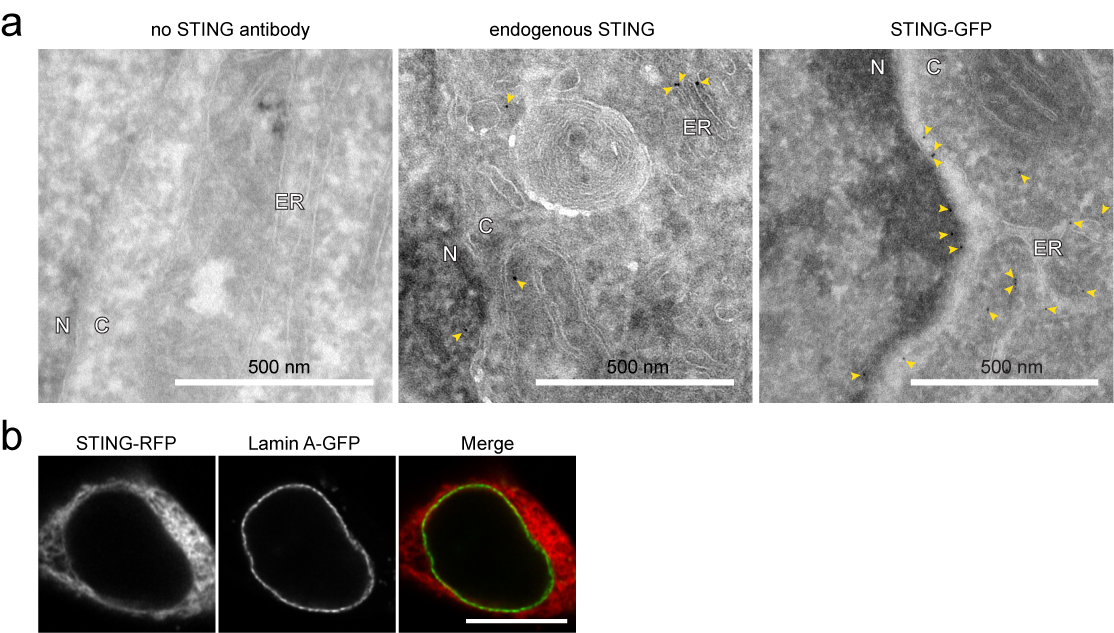

Supp Fig S2

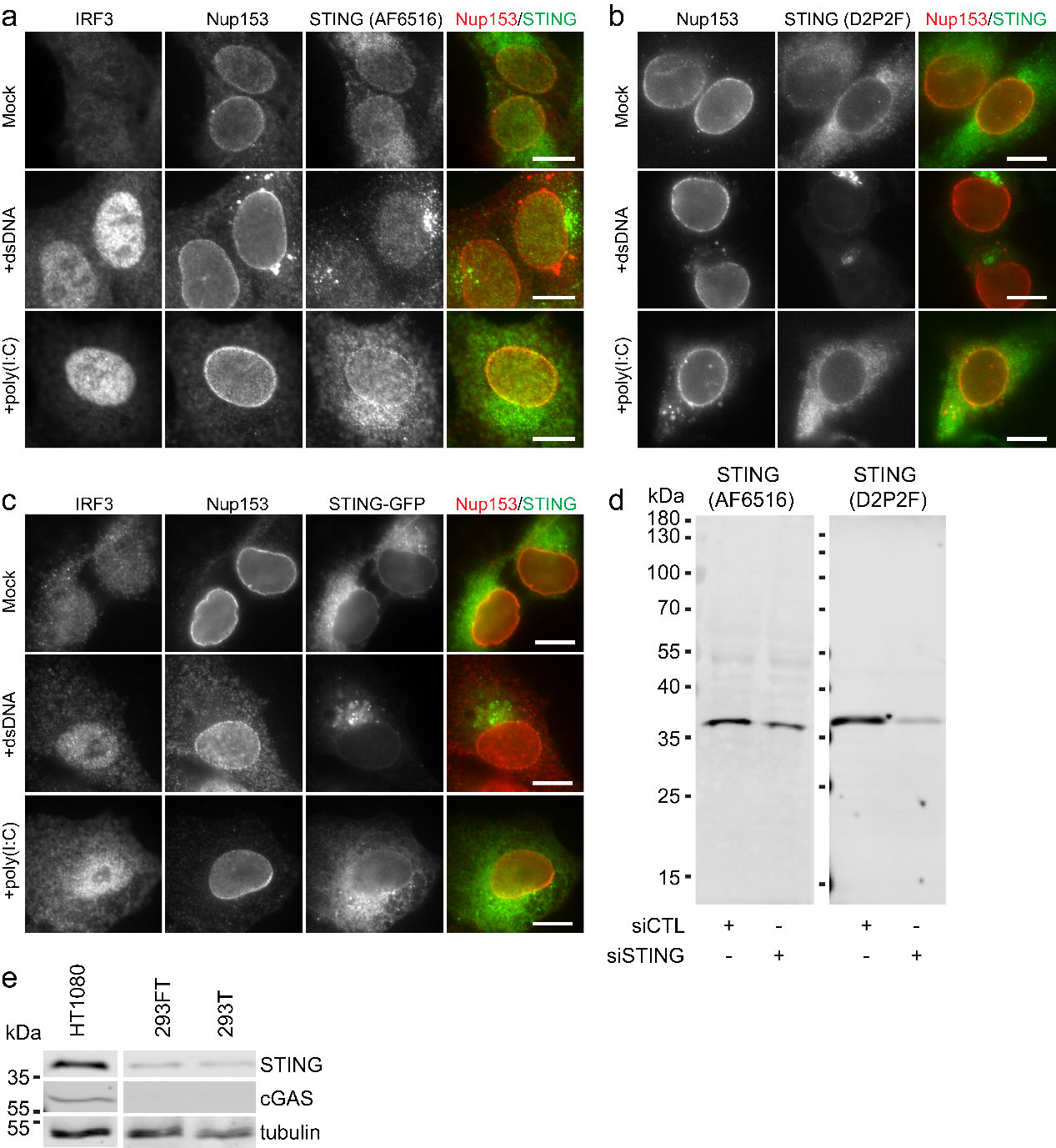

Supp Fig S3

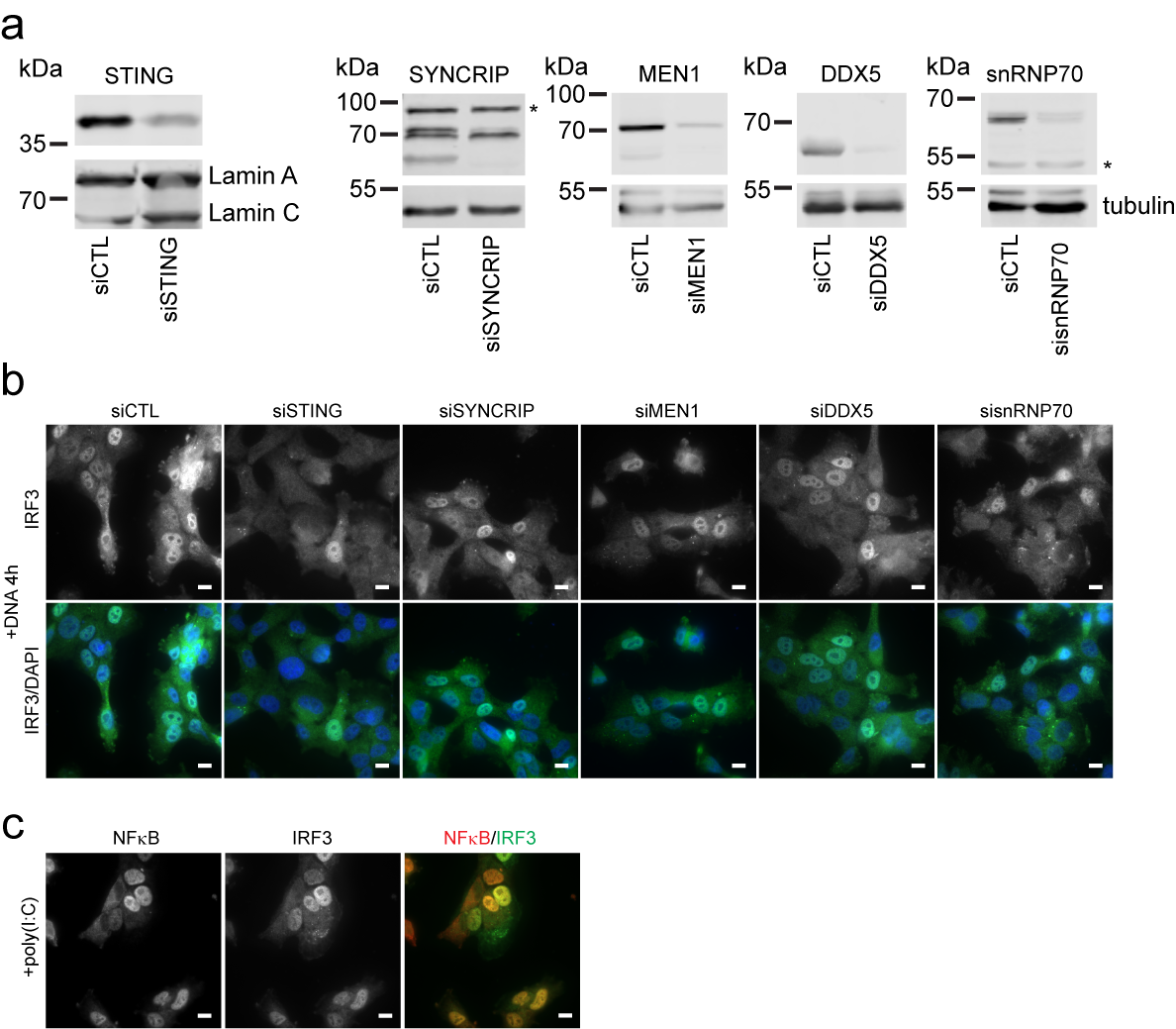
